# Impaired developmental microglial pruning of excitatory synapses on CRH-expressing hypothalamic neurons exacerbates stress responses throughout life

**DOI:** 10.1101/2021.07.21.453252

**Authors:** Jessica L. Bolton, Annabel K. Short, Shivashankar Othy, Cassandra L. Kooiker, Manlin Shao, Benjamin G. Gunn, Jaclyn Beck, Xinglong Bai, Stephanie M. Law, Julie C. Savage, Jeremy J. Lambert, Delia Belelli, Marie-Ève Tremblay, Michael D. Cahalan, Tallie Z. Baram

## Abstract

The developmental origins of stress-related mental illnesses are well-established, and early-life stress/adversity (ELA) is an important risk factor. However, it is unclear how ELA impacts the maturation of salient brain circuits, provoking enduring vulnerability to stress and stress-related disorders. Here we find that ELA increases the number and function of excitatory synapses onto stress-sensitive hypothalamic corticotropin-releasing hormone (CRH)-expressing neurons, and implicate disrupted synapse pruning by microglia as a key mechanism. Microglial process dynamics on live imaging, and engulfment of synaptic elements by microglia, were both attenuated in ELA mice, associated with deficient signaling of the microglial phagocytic receptor Mer. Accordingly, selective chemogenetic activation of ELA microglia increased microglial process dynamics and reduced excitatory synapse density to control levels. Selective early-life microglial activation also mitigated the adrenal hypertrophy and prolonged stress responses in adult ELA mice, establishing microglial actions during development as powerful contributors to experience-dependent sculpting of stress-related brain circuits.

## Introduction

Brain development is governed by both genetic factors and early-life experiences^1–4^. Specifically, adverse early-life experiences such as stress or poverty are associated with altered trajectories of structural and functional brain maturation^4,5^ and significant cognitive and emotional vulnerabilities^3,6–9^. Indeed, early-life adversity / stress (ELA) is a robust risk factor for stress-related, neuropsychiatric disorders later in life^6,7,10–12^, with profound global health implications^4,13^.

A key aspect of brain maturation involves the formation and sculpting of neuronal circuits that execute complex behaviors. This process involves axonal and dendritic growth, and especially synaptic formation, stabilization and pruning^14–19^. These take place during sensitive developmental periods and are modulated by experiences, including adversity^9,16,20–23^. However, the mechanisms by which ELA may influence and disrupt brain circuit maturation are unclear. Microglia play a key role in synaptic pruning in the developing visual and somatosensory systems^24–27^, as well as in the developing hippocampus28,29. These professional phagocytes contact and engulf synaptic elements^24,29–31^, contributing to circuit sculpting. However, a potential role for microglial dysfunction in the influence of ELA on salient brain circuit maturation has not been identified.

Here we capitalize on a robust, well-characterized model of ELA^32–34^, in which exposure to an impoverished environment and unpredictable maternal care during a sensitive developmental period provokes deficits in both emotional^35–37^ and cognitive^38,39^ functions. Importantly, in this paradigm, ELA induces enduring alteration of the responses to stress, associated with altered maturation of the hypothalamic stress-related circuit^33,40,41^. Specifically, the number of functional excitatory synapses onto stress-sensitive corticotropin-releasing hormone (CRH)-expressing neurons in the paraventricular nucleus of the hypothalamus (PVN) is augmented^41^, promoting hyperexcitability of these neurons that mediates the ELA-induced aberrant stress-responses^33,40,41^. Given the involvement of microglia in sculpting developing brain circuits elsewhere, we test here the hypothesis that ELA impairs microglia-dependent synapse pruning, resulting in augmented excitatory innervation of stress-sensitive CRH neurons and subsequent enhanced responses to stress. We find deficits in microglial process dynamics and microglial engulfment of excitatory synapses in ELA mice, which result from deficient microglial Mer receptor activity. Chemogenetic activation of ELA microglia during the developmental sensitive period restores synapse numbers and prevents the life-long augmentation of stress responses orchestrated by hypothalamic CRH neurons.

## Results

### Early-life adversity (ELA) augments the number of excitatory synapses on CRH-expressing stress-sensitive neurons

The number and function of excitatory synapses on corticotropin-releasing hormone-expressing (CRH+) neurons in the paraventricular nucleus of the hypothalamus (PVN) was augmented after a week of ELA. Specifically, the number and density of excitatory synapses, defined by the colocalization of vesicular glutamate transporter (vGlut)2 and postsynaptic density (PSD-95) per volume of CRH+ neurons, were higher in ELA pups at postnatal day (P)10 relative to mouse pups reared in control (CTL) conditions (t_12.44_=2.95, p=0.01; Welch’s t-test; Fig. 1a-b; Fig. S1a). This effect was not a result of changes in CRH+ neuron number (*p*>0.7; CTL=2475±271 cells/mm^2^; ELA= 2568±110 cell/mm^2^) or volume (*p*>0.5; CTL=31050±1773 μm^3^; ELA=32710±1741 μm^3^). These synapses endured, remaining elevated on P24-5 (t_8.96_=3.69, p=0.005; Welch’s t-test; Fig. 1c). Buttressing the neuroanatomical results, electrophysiology revealed an increased frequency of spontaneous excitatory postsynaptic currents (sEPSCs; t_19_=2.39, p=0.03; unpaired t-test; Fig. 1d-e), in mediodorsal parvocellular (mpd) PVN neurons of ELA mice, with no change in amplitude (p>0.8; Table S1), in accord with prior work^41^. Notably, these changes were specific to excitatory synapses, because neither the frequency nor amplitude of inhibitory postsynaptic currents (sIPSCs; *p*’s>0.8; Fig. 1d, f; Table S1) were altered. Together, these data are consistent with an increased number of excitatory inputs onto CRH+ neurons, resulting in an augmented excitatory drive in the PVN after ELA.

**Figure 1:**
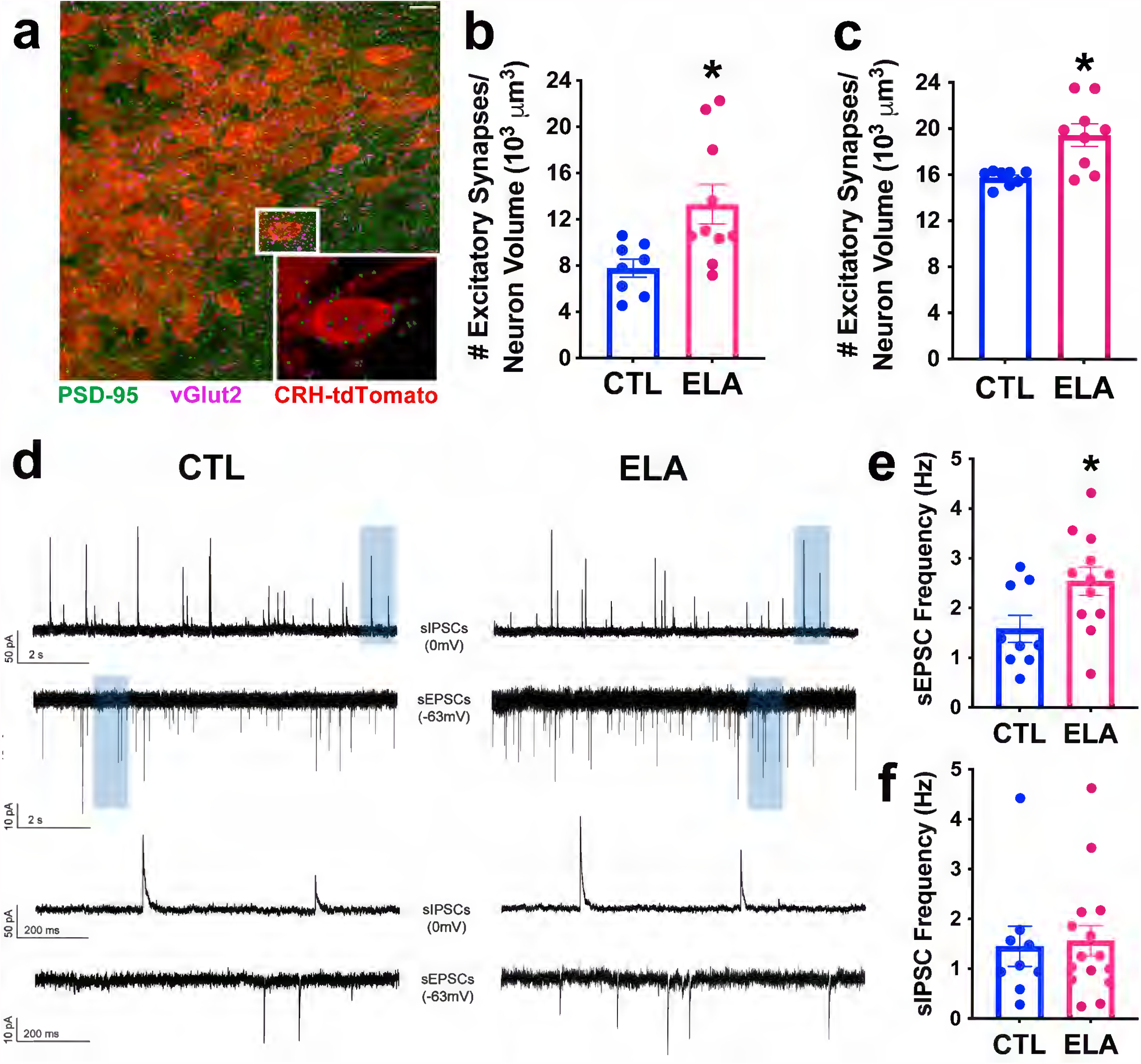
Early-life adversity (ELA) augments the number and function of excitatory synapses on CRH-expressing stress-sensitive hypothalamic neurons. a. A representative confocal image of excitatory synaptic puncta and CRH-tdTomato+ neurons in the mediodorsal parvocellular (mpd) paraventricular hypothalamic nucleus (PVN). Inset shows puncta colocalizing vGlut2-647 and PSD95-488 on a CRH-tdTomato+ neuron. These puncta satisfied criteria for synapses (Imaris software, Bitplane, Zurich, Switzerland). Scale bar=10 µm. b. ELA increases the number of excitatory synapses onto CRH+ neurons in the PVN at postnatal day (P)10 (t_12.44_=2.95, p=0.01; Welch’s t-test). c. The ELA-induced increase in excitatory synapse number endures on neurons from mice aged 24-25 days (t_8.96_=3.69, p=0.005; Welch’s t-test). d. ELA leads to functional changes in excitatory synapses of presumed CRH-expressing neurons: Representative traces are epochs (10 s) of whole-cell voltage-clamp recordings of spontaneous inhibitory synaptic currents (sIPSCs; top) and spontaneous excitatory synaptic currents (sEPSCs; bottom) recorded from mpd PVN neurons derived from CTL (left) and ELA (right) mice. A subsection (blue shaded area; 1 s) of the recorded epoch is displayed on an expanded time scale below. Scale bars: sIPSC: y = 50pA, sEPSC: y = 10pA, x = 2 seconds and 200ms for top and bottom traces, respectively. e. ELA increases the frequency of sEPSCs in mpd PVN neurons (e; t_14_=2.28, p=0.04; unpaired t-test). f. ELA does not alter the frequency of sIPSCs; (*p*>0.8). Data are mean ± SEM; \**p*<0.05.

### Process dynamics of microglia abutting PVN-CRH cells are impaired by ELA

Microglia contribute to pruning of synapses in the developing brain, a prerequisite for the formation of stable, functional brain circuits^24,28,42^. To determine whether microglial impairment underlies the observed increase of excitatory synapse density in ELA mice, we first tested for the presence of these cells in the proximity of CRH+ neurons in the developing mouse PVN. Microglia were abundant already at P4 (near the onset of the differential rearing epoch; Fig. S2a), and their neuroanatomical interactions with CRH+ neurons increased significantly during the first week of life (t_13_=3.71, p=0.003; unpaired t-test; Fig. S2b). ELA did not change the number or density of microglia abutting CRH+ cells within the PVN (p>0.1; Fig. 2a-b), nor their apparent morphology/shape. Therefore, we next interrogated microglial surveillance and phagocytic functions in ELA and control during the differential rearing period (P8). To assess microglial process dynamics, we utilized 2-photon live imaging of acute slices of PVN from dual-reporter mice (CRH-Cre+/-::tdTomato+/-; CX3CR1-GFP+/-) enabling visualization of both CRH+ neurons and microglia (Fig. 2c), and compared microglial dynamics in slices from ELA and CTL mice. We employed two independent quantification methods of microglial process excursions onto neighboring CRH+ neurons: The first, a manual kymograph method^43^ tracked the movement of the tip of a microglial process (Fig. 2d), and demonstrated that ELA significantly attenuated microglial process dynamics, as measured by the total distance traveled by process tips over 30 min. (t_16_=2.79, p=0.01; unpaired t-test; Fig. 2e). The second, a novel, automated Python-based algorithm tracked multiple microglia and their processes simultaneously (Fig. 2f) and established that ELA significantly decreased microglial process velocity (t_12_=2.26, p=0.04; unpaired t-test; Fig. 2g). Together, these independent and convergent findings suggest that microglial process dynamics, a reliable indicator of microglial function^29,44–47^, was impaired in the PVN of ELA mice.

**Figure 2.**
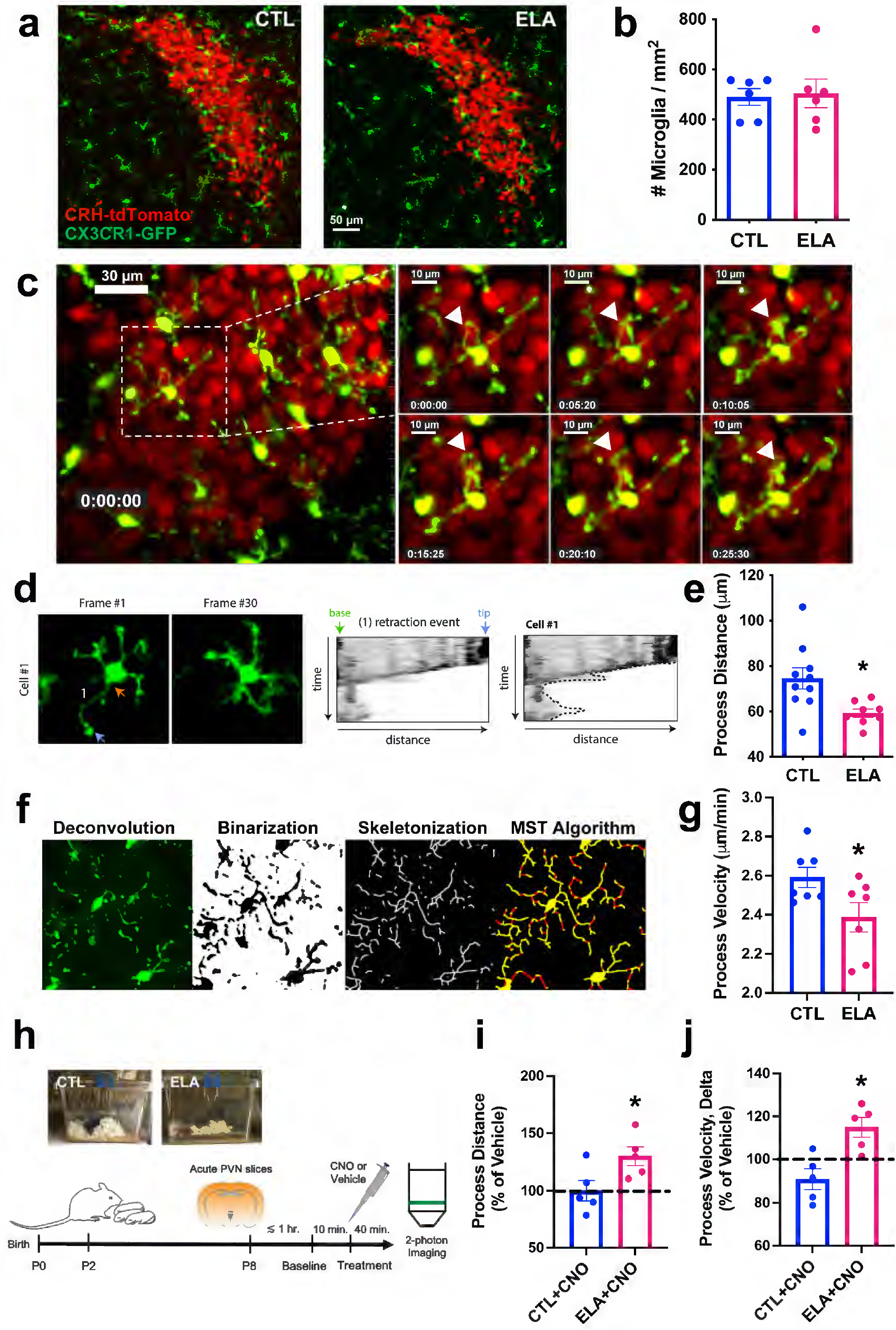
Process dynamics of microglia abutting CRH-expressing neurons in the paraventricular hypothalamic nucleus (PVN) are impaired by early-life adversity (ELA). a. Representative confocal images of microglia (green; CX3CR1-GFP) and CRH+ neurons (red; CRH-tdTomato) in the PVN of postnatal day (P)10 CTL (left) and ELA (right) mice. Scale bar=50 µm. b. ELA does not alter the density of microglia (# per region of interest [ROI] area, mm^2^) in the PVN. c. Still frames in ∼5-min. increments from a representative 2-photon video of microglia interacting with CRH+ neurons in the P8 mediodorsal parvocellular (mpd) PVN. White arrowhead points out a microglial process, highlighting that its position shifts across frames of the video. Scale bar=30 µm in leftmost image; 10 µm in smaller insets. d. Example kymograph analysis of a representative microglia from a 2-photon imaging video, tracking the retraction of a microglial process over time. In frame 1, the orange arrow points to the base of the microglial process, the location of its connection to the cell body, and the blue arrow points to the process tip. The dotted line around the edges of the rightmost kymograph represents the total distance traveled by the tip of the microglial process. e. ELA significantly attenuates microglial process dynamics, as measured by the total distance traveled by process tips over 30 min. (t_16_=2.79, p=0.01; unpaired t-test). f. Sequential steps by which microglial images in real time are processed using an automated Python-based analysis algorithm that tracks the movement of all process tips from multiple microglia simultaneously. Steps shown from left to right include: deconvolution, binarization, skeletonization, and the minimum spanning tree (MST) algorithm to fill any gaps (red) in skeletons (yellow). g. ELA reduces microglial process velocity as measured by this novel technique (t_12_=2.26, p=0.04; unpaired t-test). h. Schematic of the chemogenetic activation experiment: Litters of CX3CR1-Cre+::Gq-DREADD+ mice, born (P0), are randomly assigned to CTL or ELA rearing conditions on P2. On P8, acute hypothalamic slices are prepared, recover for ∼1 hr, and are then employed for 2-photon live imaging in superfusion chambers. Slices are imaged for a 10-min. baseline period, then exposed to CNO (to activate microglia-specific Gq-DREADDs) or vehicle in the imaging medium for a period of 40 min. i. Addition of CNO to microglia expressing excitatory Gq-DREADDs significantly augments the distance traveled by process tips *ex vivo* in ELA slices relative to vehicle treatment (t_4_=3.63, p=0.02; one-sample t*-*test) and to CTL slices (t_8_=2.45, p=0.04; unpaired t*-*test). Slices were sampled at the 30 min. time point following drug administration j. In slices from ELA mice, CNO significantly increases microglial process velocity measured at the onset and at the end of the incubation period This increase is apparent compared to vehicle treatment (t_4_=3.23, p=0.03; one-sample t*-*test) and to CTL slices (t_8_=3.64, p=0.007; unpaired t*-*test). Data are mean ± SEM; \**p*<0.05.

Aberrant microglial function is important, because their selective pruning of synapses during sensitive developmental periods may be required for the formation of refined, functional brain circuits. Therefore, we examined whether chemogenetic activation of microglia reversed the ELA-induced impairment in microglial process dynamics. We prepared acute hypothalamic slices from P8 mice expressing excitatory Gq-DREADDs specifically in microglia (CX3CR1-Cre+::Gq-DREADD+), superfused CNO or vehicle, and employed 2-photon imaging to quantify process dynamics (Fig. 2h). The addition of CNO to microglia expressing the excitatory (Gq)- DREADD significantly augmented the distance traveled by process tips *ex vivo* in ELA slices relative to vehicle treatment (t_4_=3.63, p=0.02; one-sample t*-*test; Fig. 2i) and CTL slices (t_8_=2.45, p=0.04; unpaired t*-*test; Fig. 2i) by 30 min. after the drug was added. Similarly, CNO significantly increased microglial process velocity from the beginning to the end of the drug incubation period in ELA slices relative to vehicle treatment (t_4_=3.23, p=0.03; one-sample t*-*test; Fig. 2j) and CTL slices (t_8_=3.64, p=0.007; unpaired t*-*test; Fig. 2j). Together, these findings indicate that ELA leads to reduced microglial process dynamics, and this effect cannot be attributed to non-specific toxicity because it was reversed using chemogenetic activation.

### Synapse engulfment is impaired in microglia interacting with CRH+ cells of ELA-experiencing mice

Whereas reduced microglial process dynamics has been associated with reduced “surveillance” of neurons and blunted microglial function^48–50^, it remains unclear whether these deficits in ELA microglia result in deficient synapse engulfment and pruning. We employed confocal microscopy and scanning electron microscopy (SEM) to visualize microglial engulfment of excitatory synaptic elements within the PVN (Fig. 3a-b). We first identified P8 as a period of active synapse engulfment, establishing a significantly higher number and density of engulfed vGlut2+ synaptic puncta in microglial elements on P8 compared with P30 in CTL mice (Fig. S3a). Focusing on P8, we identified in ELA mice fewer vGlut2+ synaptic puncta engulfed by microglia compared with CTLs (t_13.87_=2.22, *p*=0.04; Welch’s t-test; Fig. 3c). We excluded the possibility that this reduction resulted from ELA-induced increases in the total volume occupied by microglia in the PVN (*p*>0.1; CTL=12754±674 μm^3^; ELA=14079±705 μm^3^. Reduced synapse engulfment was selective to microglia abutting CRH+ neurons; vGlut2+ synapse engulfment in microglia on the periphery of the PVN that were not in direct contact with CRH+ neurons did not differ in CTL and ELA mice (*p*>0.2; Fig. S3b). Together, these data suggest that reduced process dynamics associates with, and likely contributes to, poor synapse engulfment in microglia abutting CRH+ neurons in ELA mice.

**Figure 3:**
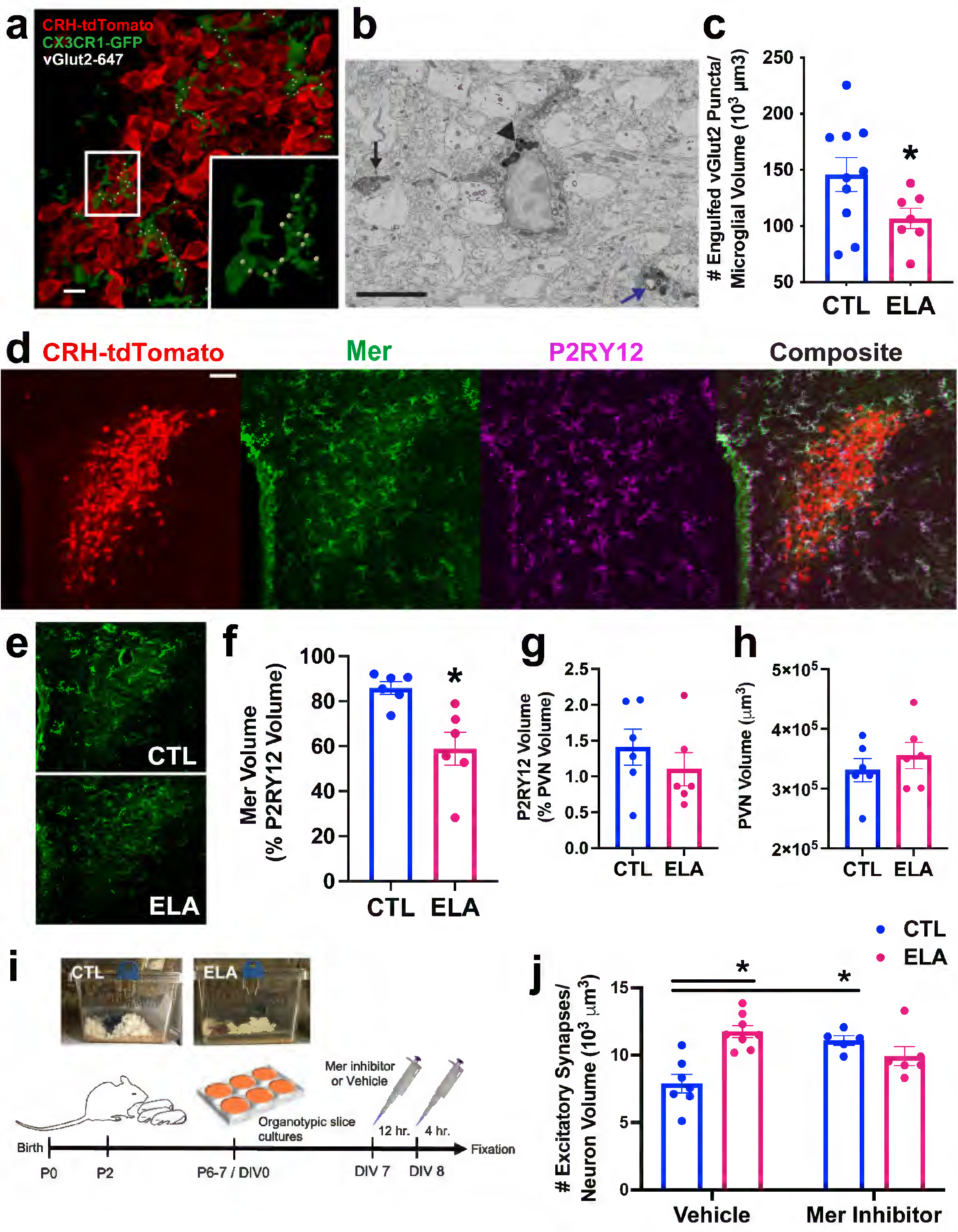
Microglial mechanisms for augmented excitatory synapses on CRH+ cells in ELA mice. a. Representative confocal image of microglia (green; CX3CR1-GFP) abutting CRH+ neurons (red; CRH-tdTomato) and engulfing vGlut2+ excitatory presynaptic puncta (white; Imaris 3D reconstruction software) in the postnatal day (P)8 mediodorsal parvocellular (mpd) paraventricular hypothalamic nucleus (PVN). Scale bar=10 µm. b. Representative electron micrograph of a microglia labeled with ionized calcium binding adaptor molecule (Iba)1 and 3’3-diaminobenzidine (DAB; darker, electron-dense cell body and processes) in the P8 PVN. The black arrow points to the postsynaptic density of an excitatory synapse directly abutting a labeled microglial process. The black arrowhead indicates multiple lysosomes (electron-dense) in the microglial cell body; these degrade engulfed material. The blue arrow points to an engulfed endosome of synaptic elements, including a postsynaptic density, surrounded by lysosomes in the microglial process. Scale bar=5 µm. c. ELA reduces the number of vGlut2+ synaptic puncta engulfed by microglia abutting CRH+ neurons in the P8 mpd PVN (t_13.87_=2.22, *p*=0.04; Welch’s t-test; Fig. 3c). d. Representative confocal image of the microglial phagocytic receptor Mer tyrosine kinase (Mer) expression. The receptor is localized primarily in microglia in the P8 PVN. Red= CRH-tdTomato+ neurons; green= Mer immunoreactivity; magenta= P2RY12 immunoreactivity (labels microglia); white= overlap of green and magenta in composite image. Scale bar=50 µm. e. Representative confocal images of Mer immunoreactivity in CTL (top) vs. ELA (bottom) P8 PVN. f. Mer levels per unit volume of P2RY12+ microglia (measured using Imaris software) were reduced in PVN of P8 ELA mice compared with controls (t_6.46_=3.44, *p*=0.01; Welch’s t*-*test). g. Notably, P2RY12 volume, as a percentage of PVN ROI volume, did not differ in ELA and control mice. h. PVN volume also did not distinguish ELA and CTL mice. i. Schematic of the Mer inhibition experiment: Litters of CRH-tdTomato+ mice were born (P0) and randomly assigned to CTL or ELA rearing conditions on P2. On P6-7, organotypic hypothalamic slice cultures were prepared and maintained for 7 days in vitro (DIV). On DIV7, cultures were treated with 20 nM of a small-molecule Mer inhibitor or vehicle. 12 hr. later, media was refreshed with new drug, and incubated for 4 more hr. until fixation. j. Mer inhibition increased the number of excitatory synapses on CRH+ neurons in PVN cultures from control mice but failed to augment synapse number in PVN cultures of ELA mice (significant interaction of ELA x Drug, F_1,22_=17.89, p=0.0003; 2-way ANOVA; p<0.05; post hoc Tukey’s test). Data are mean ± SEM; \**p*<0.05.

### Mechanisms of impaired synapse pruning in ELA microglia involve the phagocytic receptor Mer

Microglia prune synapses in response to several cellular and molecular triggers. Among them, the microglial phagocytic receptor, Mer tyrosine kinase^48,51,52^ has been reported in hippocampus. Therefore, we tested whether this receptor contributed to the change in synaptic number in ELA mice. We first established Mer expression in the P8 PVN and localized it predominantly to microglia (Fig. 3d). There was no overlap of Mer with the astrocytic markers GFAP and S100β or the neuronal marker NeuN; (Fig. S4). The presence of Mer selectively in hypothalamic microglia during the developmental time point of interest prompted assessment of Mer levels in the PVN of ELA and control mice. Indeed, at P8, Mer immunoreactivity relative to microglial volume was diminished in the PVN of ELA mice (Fig. 3e) (t_6.46_=3.44, *p*=0.01; Welch’s t*-*test; Fig. 3f). Importantly, neither P2RY12 immunoreactivity nor PVN volume were altered in ELA mice (*p*’s>0.3; Fig. 3g-h). We then tested if deficient Mer signaling might explain the attenuated synapse engulfment by ELA microglia. This hypothesis predicts that blocking Mer would increase synapse number in control but not in ELA-derived CRH+ hypothalamic neurons, in which the receptor is already less active. To test this prediction, we employed hypothalamic organotypic slice cultures containing the PVN and treated them with a small-molecule Mer inhibitor or vehicle for 16 hr. (Fig. 3i). Mer inhibition increased the number of excitatory synapses on CRH+ neurons in PVNs from control mice but failed to augment synapse number in PVNs of ELA mice (significant interaction of ELA x Drug, F_1,22_=17.89, p=0.0003; 2-way ANOVA; p<0.05, post hoc Tukey’s test; Fig. 3j). These data support the notion that microglial Mer signaling is already inhibited in ELA mice, and underlies the increase in excitatory synapses.

### Functional significance of ELA-induced microglial deficits

To determine whether the ELA-induced microglial deficits during development are functionally significant for long-lasting stress-related outcomes governed by hypothalamic CRH+ cells, we chemogenetically activated microglia in developing CTL and ELA mice and determined the hormonal responses of these mice to acute stress during adulthood. We also assessed the overall life-time neuroendocrine responses of these mice to stress by measuring the weight of the adrenal glands, as chronic augmented stress-responses lead to enlarged, heavier glands^53^. For selective chemogenetic activation of microglia during a defined early-life period, we crossed CX3CR1-Cre+::Gq-DREADD+ mice with Gq-DREADD+ mice. This results in litters in which ∼50% of pups express Gq-DREADDs in their microglia (Fig. 4a; Fig. S5) and the rest serve as littermate controls. We inserted small, sustained-release CNO or placebo-containing pellets under the skin (s.c.) on P3 and randomized the mice to CTL or ELA rearing for a week (Fig. 4b).

**Fig 4:**
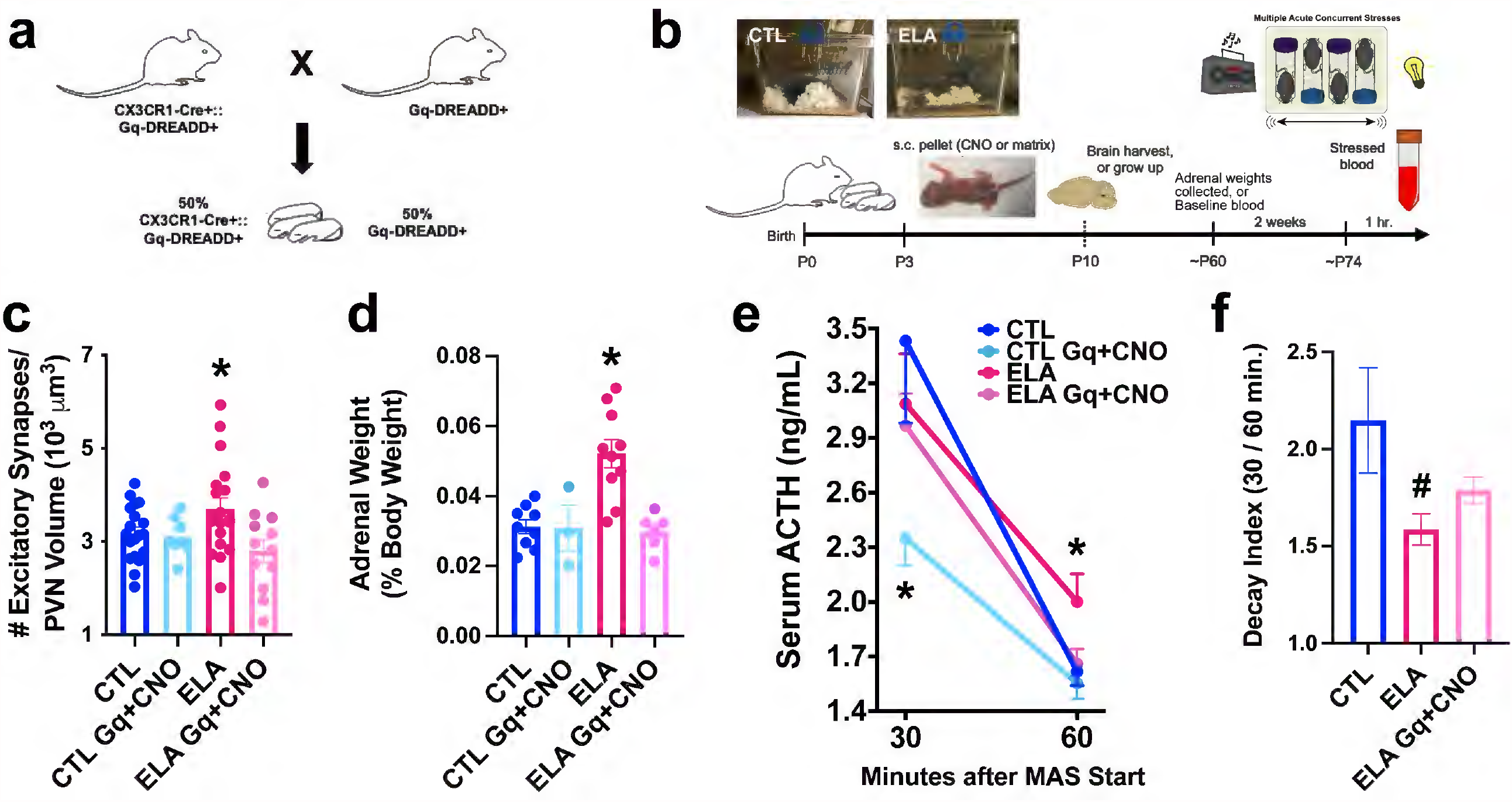
Enduring functional significance of ELA-related microglial deficits in synapse pruning. a. Breeding strategy for the chemogenetic studies: CX3CR1-Cre+::Gq-DREADD+ mice were crossed with Gq-DREADD+ mice to generate ∼50% pups expressing Gq-DREADDs exclusively in microglia. b. Schematic of *in vivo* chemogenetic activation experiment: Litters of CX3CR1-Cre+::Gq-DREADD+ pups, born (postnatal day [P]0) and randomly assigned to CTL or ELA rearing conditions (P3) and received small, sustained-release CNO- or placebo-containing pellets under the skin (s.c.). One cohort of these mice was perfused on P10 for quantification of excitatory synapses (colocalized vGlut2+PSD95) onto mediodorsal parvocellular (mpd) paraventricular hypothalamic nucleus (PVN) cells. A separate cohort of mice provided adrenal gland weights, a measure of lifetime stress responses as adults. A third cohort provided baseline and stress-induced ACTH and corticosterone (CORT) levels. As adults, these mice experienced an acute multimodal stress (MAS60) and blood was collected at 30 and 60 min. following stress onset. c. Chronic chemogenetic microglial activation during postnatal days 3-10 in microglia-specific Gq-DREADDs in ELA mice decreased the number of excitatory synapses on mpd PVN neurons to control levels; this was not observed in ELA mice lacking microglial expression of Gq-DREADDs or in placebo-receiving mice (F_2,23.8_=3.76, p=0.04; Welch’s one-way ANOVA; *p*<0.05, Dunnett’s T3 multiple comparisons test). Because CNO treatment alone did not alter the number of excitatory synapses in microglial Gq-DREADD-expressing CTL mice (*p*>0.6), CTL groups were combined for analysis. d. Adrenal weights of ELA-experiencing adult mice were higher than those of control mice, indicative of lifelong exposure to augmented stress-hormone release. Remarkably, chemogenetic microglial activation during the ELA epoch prevented the adrenal weight increase (F_3,24_=11.11, p<0.0001; one-way ANOVA; *p*<0.05, Holm-Sidak’s multiple comparisons test). e. At 30 min. from onset, MAS elicited a robust elevation of the stress hormone ACTH in both CTL and ELA mice. Chemogenetic activation of microglia in CTL mice blunted this response (F_3,20.6_=3.19, p=0.04; Welch’s one-way ANOVA; p<0.05, Dunnett’s T3 multiple comparisons test). At 60 min. after stress onset, ACTH levels dropped to 50% of peak values in CTL mice, but not in ELA mice (F_2,53_=3.45, p=0.04; one-way ANOVA; *p*<0.05, Holm-Sidak’s multiple comparisons test). Chemogenetic microglial activation in ELA mice ameliorated this prolonged elevation of ACTH levels, which returned to control levels by 60 min. (*p*>0.8, Holm-Sidak’s multiple comparisons test). f. The ACTH decay index (30 min./60 min. ACTH levels) differed in ELA mice (F_3,21.5_=3.19, *p*=0.04; Welch’s one-way ANOVA; *p*=0.06, Dunnett’s T3 multiple comparisons test), and was partially restored in ELA mice whose microglia were chemogenetically activated (*p*>0.2 vs. CTL, Dunnett’s T3 multiple comparisons test). These data indicate an aberrant prolongation of the neuroendocrine stress response in ELA mice, and the restoration of normal ‘shut-off’ mechanisms by early-life microglial activation. (CTL Gq+CNO mice were not included in the decay index analysis because their stress response was already blunted.) Data are mean ± SEM; \**p*<0.05.

Outcome measures included the number of excitatory synapses onto PVN mpd neurons at the end of the week (P10). In separate cohorts we measured levels of ACTH and corticosterone (CORT) that depend on CRH release from hypothalamic CRH cells^54,55^ in response to an acute stress, as well as adrenal weights, both during adulthood (>P60).

In ELA mice expressing microglia-specific Gq-DREADDs, CNO reduced the number of excitatory synapses on presumed CRH cells in PVN to control levels. CNO had no effect in ELA mice lacking microglial expression of Gq-DREADDs, and placebo-containing pellets also did not alter synapse numbers (F_2,23.8_=3.76, p=0.04; Welch’s one-way ANOVA; *p*<0.05, Dunnett’s T3 multiple comparisons test; Fig. 4c). Of note, CNO treatment did not alter the number of excitatory synapses in microglial Gq-DREADD-expressing CTL mice (*p*>0.6), so that all the CTL groups were combined for analysis. These data indicate that artificial activation of microglia during a sensitive developmental period prevents the increase in synapse number provoked by ELA, supporting the notion that impaired microglial function resulting from ELA is the mechanism for the increased excitatory synapses on CRH neurons in the PVN.

We next determined the enduring consequences of ELA and of activating microglia early in life on the hormonal responses of adult ELA mice to stress, and measured adrenal weights as a proxy for lifelong, chronic stress levels^53,56,57^ in adult CX3CR1-Cre+::Gq-DREADD+ given CNO or vehicle early in life. ELA augmented adrenal weights, in accord with our prior work^38,58^. Remarkably, chemogenetic microglial activation prevented ELA-induced adrenal weight increase (F_3,24_=11.11, p<0.0001; one-way ANOVA; *p*<0.05, Holm-Sidak’s multiple comparisons test; Fig. 4d). Complex, acute stress^59,60^ elicits a robust elevation of the stress hormones adrenocorticotropic hormone (ACTH) and corticosterone (CORT), which is governed by CRH+ hypothalamic neurons. In response to this stress, significant elevations of plasma ACTH and corticosterone were observed in both control and ELA-reared adult mice, and did not differ at 30 min. (ACTH, Fig. 4e, and CORT, not shown) or 60 min. (CORT, not shown) after stress onset. However, the decline kinetics of ACTH, or termination of the hormonal response to stress, differed significantly among groups: At 60 min. after stress onset, ACTH levels were 50% lower than their peak in CTL mice; by contrast, the decline of ACTH levels was attenuated in ELA mice (F_2,53_=3.45, p=0.04; one-way ANOVA; *p*<0.05, Holm-Sidak’s multiple comparisons test; Fig 4e). This aberrant prolongation of the stress response was also apparent as a lower ACTH decay index (30 min./60 min. F_3,21.5_=3.19, *p*=0.04; Welch’s one-way ANOVA; *p*=0.06, Dunnett’s T3 multiple comparisons test; Fig. 4f) in ELA mice, in accord with our prior studies demonstrating an abnormally lengthy stress response after ELA^33^. Strikingly, early-life chemogenetic microglial activation in ELA mice prevented the persistent elevation of ACTH levels in adulthood: ACTH levels at 60 min. were similar to those in CTL mice (*p*>0.8, Holm-Sidak’s multiple comparisons test; Fig. 4e), accompanied by partial restoration of the decay index (*p*>0.2 vs. CTL, Dunnett’s T3 multiple comparisons test; Fig. 4f). Together, these results demonstrate that ELA-induced microglial dysfunction in developing PVN has important, enduring consequences on acute and chronic responses to stress later in life.

## Conclusions

The principal findings described here are: 1) In the developing stress circuit, microglia are critical for normal synapse pruning and the maturation of structural and functional connectivity of stress-sensitive CRH+ neurons; 2) microglial process dynamics may be a potent indicator and a potential mechanism of their synapse pruning capacity during development; 3) ELA provokes microglial dysfunction during development, which results in augmented excitatory synapses on CRH+ neurons in the PVN and consequent enhanced reactivity to future stress; 4) the mechanism for impaired synapse engulfment during ELA involves differential expression and activity of the microglial phagocytic receptor Mer. These discoveries identify an important role for microglia actions during a sensitive period in ELA-induced enduring stress sensitivity, a hallmark of stress-related affective disorders.

These studies demonstrate that microglia are critical for the normal, selective synapse pruning during a sensitive developmental period in a stress-related circuit, and specifically in a subpopulation of stress-sensitive CRH+ neurons in the PVN. Previous work has demonstrated that microglia engulf synapses in other developing brain regions, including the thalamus^24^, hippocampus^28^, and cortex^25,61^. However, prior analyses did not consider cell-type specificity of the neuronal targets of microglial synapse pruning. Recent data have supported the notion that microglia are a heterogenous population of cells^62,63^, differing across brain regions and even within a brain region^62,64^. Here, we demonstrate that as a result of ELA, microglia engulf fewer synaptic elements of CRH+ neurons in the PVN (Fig. 3), whereas the synapse engulfment of microglia not in contact with CRH+ neurons was not affected (Fig. S3). These results suggest a selectivity of the neuronal targets of microglia, at least within the PVN, which may derive from altered signaling of CRH cells to adjacent microglia as a result of ELA^65,66^. Alternatively, distinct microglia populations may be assigned to synapse pruning of specific neuronal populations within the PVN. Distinguishing between these two possibilities will require future studies.

Notably, microglial synapse pruning of PVN-CRH+ neurons influences the function of the neuroendocrine responses to stress, i.e., the hypothalamic-pituitary-adrenal (HPA) axis, because the augmented number of excitatory synapses observed after ELA results in increased excitatory signaling in the PVN (Fig. 1), as well as higher adrenal weights, indicative of lifelong chronic stress, and prolonged secretion of stress hormones in response to an adult stressor (Fig. 4). The specific role of microglia in these synaptic and functional changes is evident from our demonstration that selective chemogenetic activation of microglia *in vivo* during a sensitive period prevents ELA-induced augmentation of excitatory synapse number and function, and restores adult stress sensitivity to control levels (Fig. 4). Together, these data demonstrate a causal link between microglial function during development and lifelong stress reactivity.

We quantified here microglial process dynamics in real time. Conventional measures of microglial activation, such as microglial density and shape, provide limited insight into microglial function during development when these measures are in flux^67^. We did not find differences in microglial density or shape in ELA hypothalamus and instead obtained direct measures of microglial function using two salient, independent parameters: microglial process dynamics and synapse engulfment. ELA inhibited both of these measures of microglial function, suggesting that they may be related: process dynamics such as velocity and excursion distance may predict and perhaps underlie synapse engulfment efficiency. Thus, deficits in microglial process dynamics, and potentially actin cytoskeleton remodeling, could result in diminished synaptic engulfment. Alternatively, deficits in microglial synapse pruning, mediated via Mer and/or its ligand expressed on neurons, could reduce surveillance of the extracellular environment by microglial processes. Future studies should test the validity of these two hypotheses, and thus offer further insight into the true nature of microglial function and activation during development.

Mechanistically, we discovered that the regulation of excitatory synapse number on CRH+ neurons by microglia within the PVN was dependent on the phagocytic receptor Mer. *Ex vivo* treatment of PVN organotypic slices with a small-molecule Mer inhibitor augmented excitatory synapses on CRH+ neurons (Fig. 3), indicating that microglial synapse pruning was at least partially inhibited. Mer is expressed by microglia^51,52^ and astrocytes^68^ in the brain. In the immature PVN, we localized Mer expression exclusively to microglia (Fig. 3 and Fig. S4). Recent work implicates astrocytes in synaptic pruning^68,69^, likely in concert with microglia^19,30^, and these interactions merit future studies. Here, ELA reduced Mer expression in microglia of the developing PVN. In addition, whereas Mer inhibition increased synapse numbers in control PVN cultures, it did not influence synapse numbers in in PVN cultures from ELA mice (Fig. 3), suggesting that this pathway was already attenuated in ELA mice. Mer interacts with a neuronal ligand, phosphatidylserine, an intracellular molecule that is externalized during cell death ^51,52^, providing a phagocytic signal. Recently, selective Mer externalization at synaptic elements has been identified, and implicated in developmental synaptic pruning^70^. It is not known if externalization of phosphatidylserine at synapses on CRH+ neurons is reduced by ELA, attenuating the signal for microglial synaptic pruning. The data presented here suggest that expression and function of Mer is inhibited in ELA hypothalamus. This, potentially in concert with diminished externalization of its ligand, results in the diminished microglial synapse pruning observed in the PVN during ELA.

How might ELA signal to microglia? ELA is associated with chronic elevation of plasma glucocorticoids^33,38,40^, and glucocorticoids reduce microglial function^71–73^. In addition, the stress hormone CRH may be released also within the PVN during ELA. Microglia express both glucocorticoid receptors and CRFR1^74^ and may be directly affected by these hormones. Alternatively, they may be impacted indirectly through activity-dependent neuronal signals, such as ATP or adenosine^44,45,73^. Our ability to rescue microglial function chemogenetically in ELA mice suggests the effect of ELA on these cells can be overcome by interventions delivered directly to microglia, holding promise for future therapeutic strategies.

In summary, the findings described here establish microglia as powerful contributors to enduring, experience-dependent sculpting of stress-related brain circuits. ELA provokes microglial dysfunction during a sensitive period of development, characterized by diminished microglial process dynamics and deficits in microglial synapse engulfment, and resulting in augmented number and function of excitatory synapses on CRH+ neurons in the PVN. These changes are critical, because they govern lifelong chronic stress hormone levels and stress reactivity, an important factor in determining risk for stress-related mental illnesses. The manipulation of microglial function during development may provide novel targets for prevention of stress-related disorders.

## Acknowledgments

This work was supported by NIH grants K99 MH120327 (JLB), P50 MH096889 (TZB), R01 MH73136 (TZB), R01 NS14609 (MDC), and R01 AI121945 (MDC); the Brain & Behavior Research Foundation NARSAD Young Investigator Grant (JLB), and the Hewitt Foundation for Biomedical Research (TZB). We thank the Optical Biology Core Facility of the Developmental Biology Center, supported by the Cancer Center Support Grant (CA-62203) and Center for Complex Biological Systems Support Grant (GM-076516) at the University of California-Irvine. We thank Dr. Ian Parker, UCI for access to the custom-built 2-photon microscope used for the live imaging. The CX3CR1-BAC-Cre mice were kindly provided by Dr. Staci Bilbo (Duke University). We thank Emily Majorkiewicz, Andrew Quan Dong, Xinwen Li, Yanan Wu, Catherine Chiou, Keshav Suresh, Kathleen Guangying Zhou, Pouria Vadipour, Graciella Angeles, Johnathan Ho, Erin Card, and Jennifer Daglian for excellent technical assistance. We also thank Dr. Yoav Noam for his insight and support with kymograph analyses.

## Author Contributions

JLB and TZB designed the experiments. JLB, MS, SO, SML, AKS, CLK, BGG, and JCS performed the experiments. JB and XB developed analytical tools. JLB, MS, SO, SML, AKS, JB, XB, CLK, BGG, and TZB analyzed data. JJL, DB, MET and MDC provided conceptual advice, access to critical equipment, and supervised collaborative work in their respective laboratories. JLB and TZB wrote and/or edited the paper.

## Methods

### Animals

Mice of both sexes were housed in temperature-controlled, quiet, uncrowded conditions on a 12-hr light, 12-hr dark schedule (lights on at 0630 hr., lights off at 1830 hr.) with free access to food and water. The transgenic mouse lines employed were all on a C57BL/6 genetic background bred in-house from the following founders: CRH-IRES-Cre+/+ (Jax #: 012704; Jackson Laboratory, Bar Harbor, ME), tdTomato+/+ (stop-floxed; Ai14; Jax #: 007914), CX3CR1-GFP+/+ (Jax #: 005582), Gq-DREADD+/- (stop-floxed; Jax #: 026220) CX3CR1-BAC-Cre+ (GENSAT; MMRRC stock #036395-UCD; obtained from Dr. Staci Bilbo, Duke University). Wild-type C57BL/6J mice used for the electrophysiology experiments were bred in-house from founders obtained from Jax (#000664). All experiments were performed in accordance with National Institutes of Health guidelines and were approved by the University of California, Irvine Animal Care and Use Committee.

### Limited Bedding and Nesting (LBN) Model of Early-life Adversity (ELA)

Dams were checked for copulatory plugs daily while paired with a male, housed singly on embryonic day (E)17, and monitored every 12 hr. for the birth of pups. The day of birth was termed postnatal day (P)0. On the morning of P2, litters were culled to 8 pups if needed (5 pups minimum) and included both sexes. Also on P2, the ELA paradigm was initiated as in previous studies^34,40^: Control (CTL) dams were placed in cages with standard amounts of corn husk bedding (650 ml) and nesting material, i.e. one square piece of cotton material measuring 5 × 5 cm (nestlet). This material was shredded by the dam to create a nest area. In contrast, ELA dams received one-half nestlet placed on a fine-gauge plastic-coated aluminum mesh platform (mesh dimensions 0.4 × 0.9 cm, cat# 57398; McNichols Co., Tampa, FL), approximately 2.5 cm above the cage floor. The cage floor was covered with a small amount of corn husk bedding (∼60 ml). This setup permitted mouse droppings to fall below the platform without trapping the pups. In addition, all cages were housed in a room with robust ventilation, avoiding the accumulation of ammonia. Figure S1a shows examples of the ELA and standard cage setups. Both groups were completely undisturbed from the morning of P2 to the morning of P10, at which point the dams and litters were transferred to standard cages.

The impoverished cage environment in mice results in marked alterations in maternal behavior, such as increased sorties (i.e., exits) from the nest during P2-8 (F_1,19_=25.15, p<0.0001; 2-way repeated measures ANOVA; Fig. S1b) and shorter, fragmented bouts of licking and grooming the pups (t_18_=5.26, p<0.0001; unpaired t-test; Fig. S1c), without changes in the total amount of licking and grooming behaviors (Fig. S1d), in accord with our previous studies^35,40^. This environment provokes chronic stress in the mouse pups, including reduced weight gain on P10 (t_101_=7.05, p<0.0001; unpaired t-test; Fig. S1e) and P21 (t_41_=3.45, p=0.001; unpaired t-test; Fig. S1f). These modest changes disappear by adulthood (P60; Fig. S1g). We have also previously identified increased adrenal weights and serum corticosterone levels during the ELA period^38,40,75^.

Both sexes were used for all initial experiments (i.e., synapse counts, electrophysiology; Fig. 1, Fig. S6). These studies demonstrated an additional effect of sex. Specifically, unlike males, increased excitatory synapse number and function on CRH+ neurons in the PVN by P10 (Fig. S6a) was not observed in females. Notably, by the juvenile period (P24-5; Fig. S6b), both males and females were affected. In addition, microglial synapse engulfment at P8 was not altered by ELA in females (Fig. S6c). Because these data suggested sex-modulated time course of synapse development and contributions of ELA, we elected to focus the subsequent studies on males alone, and plan to devote future work to the studies of females.

For short-term studies, pups were euthanized with sodium pentobarbital and transcardially perfused with ice-cold phosphate-buffered saline (PBS; pH=7.4) followed by 4% paraformaldehyde in 0.1M sodium phosphate buffer (pH=7.4) on P8 (for microglial synapse engulfment studies) or P10 (for synapse counts). For 2-photon imaging studies, pups were euthanized via rapid decapitation and acute slices of the paraventricular hypothalamic nucleus (PVN) prepared. For long-term studies in the juvenile period (i.e., electrophysiology, synapse counts) and adulthood (i.e., functional studies of stress response), pups were weaned on P21 into standard, same-sex cages of 2-5 mice.

### Assessment of maternal behaviors in CTL vs. ELA cages

Maternal behaviors and interactions with pups were evaluated on P2–8. Control (n=9 litters) and ELA dams (n=11) were observed daily in the morning (between 0700-0900 hr.). Each maternal observation session consisted of 50 min, during which maternal behaviors and dam-pup interactions were scored continuously, allowing for an accurate analysis of the duration of each bout of behavior^35^. The behaviors recorded included: licking and grooming pups (LG), nursing pups, self-grooming, nest-building, off-nest activity, and eating/drinking. The number of times the dam left the nest (sorties) was also recorded^40^. To optimize visibility into the nest and minimize interference with normal cage activities, the observer used mirrors placed beside the cages.

### Immunohistochemistry (IHC) for Excitatory Synaptic Markers

Perfused brains of CRH-Cre+/-::tdTomato+/- mice (P10 or P24-5) were post-fixed in 4% paraformaldehyde in 0.1 M PBS (pH = 7.4) for 4-6 hr. before cryoprotection in a 25% sucrose solution. Brains were frozen, then sectioned coronally into 25-µm-thick slices (1:4 series of the PVN) using a Leica CM1900 cryostat (Leica Microsystems, Wetzlar, Germany). Sections were subjected to IHC using standard methods, as described previously^76^. Briefly, after several washes with PBS containing 0.3% Triton X-100 (PBS-T, pH 7.4), sections were blocked with 5% normal donkey serum (cat# 017-000-121, Jackson ImmunoResearch, West Grove, PA, USA) for 1 hr. to prevent non-specific binding. Sections were then incubated overnight at 4°C with rabbit anti-PSD95 antiserum (1:1,000; cat# 51-6900, Invitrogen/ThermoFisher, Waltham, MA) and guinea pig anti-vGlut2 antiserum (1:12,000, cat# AB2251-1, Millipore Sigma, Temecula, CA, USA) in PBS-T containing 2% normal donkey serum. The next morning, sections were rinsed in PBS-T (3 × 5 min.), and then incubated with donkey-anti-rabbit IgG-488 (1:1,000; A-21206, ThermoFisher) and donkey-anti-guinea pig IgG-647 (1:1,000; cat# 706-605-148, Jackson Immunoresearch) for 3 hr. at room temperature. After washing (3 × 5 min), sections were mounted onto gelatin-coated slides and coverslipped with Vectashield containing DAPI (cat. #H-1200, Vector Laboratories, Burlingame, CA, USA).

#### Confocal imaging

Confocal images of the mediodorsal parvocellular (mpd) paraventricular hypothalamic nucleus (PVN) were collected using an LSM-510 confocal microscope (Zeiss, Dublin, CA, USA) with an Apochromat ×63 oil objective. 11 z-stack images of 142.86 × 142.86 μm were taken at 1-μm intervals. Image frame was digitized at 12-bit using a 1024 × 1024 pixel frame size. CRH+ neuron number was counted manually in FIJI, and CRH neuronal volume was automatically calculated using Imaris’ 3D reconstruction. Excitatory synapses onto CRH+ neurons were identified as colocalized puncta of vGlut2+PSD95 within the CRH-tdTomato+ volume using Imaris’ colocalization function (threshold=1.0).

### Electrophysiology

#### Slice preparation

Wild-type control and ELA mice (P18-26) were killed by cervical dislocation in accordance with Schedule I of the UK Government Animals (Scientific Procedures) Act, 1986. Coronal hypothalamic slices containing the PVN were prepared as previously described41. The brain was rapidly dissected and placed in ice cold (0-4°C), oxygenated (95% O_2_) artificial cerebro-spinal fluid (aCSF) containing the following (in mM): 135 NaCl, 2.5 KCl, 1.25 NaH_2_PO_4_, 10 MgCl_2_, 0.5 CaCl_2_, 26 NaHCO_3_, 10 glucose (320-335 mOsm, pH ∼7.4). Coronal hypothalamic slices were then cut (300-320 μm) using a vibratome (Leica) at 0-4°C. Slices were subsequently incubated for at least 1 hour at room temperature in a holding chamber containing oxygenated aCSF (as above) that additionally contained 1 mM ascorbic acid and 3 mM sodium pyruvate. Slices were then transferred to the recording chamber and recordings were performed on slices perfused (3-6 ml/min) with extracellular solution (ECS) containing (in mM): 126 NaCl, 26 NaHCO_3_, 2.95 KCl, 1.25 NaH_2_PO_4_, 2 MgCl_2_, 2 CaCl_2_, 10 glucose (306-309 mOsm, pH ∼7.4).

#### Whole-cell recordings

All recordings were performed using an Axopatch 1D amplifier (Molecular Devices, Wokingham, UK), stored directly to a PC using a NI-DAQmx interface (National Instruments, Newbury, UK) for analysis offline.

#### Recordings of sEPSCs and sIPSCs

Whole-cell voltage-clamp recordings of sEPSCs and sIPSCs were made at 35°C in ECS using a low Cl- (12 mM) intracellular solution. In these recordings, patch pipettes (4-6 MΩ) were filled with an intracellular solution containing (in mM): 135 CH_3_O_3_SCs, 8 CsCl, 10 HEPES, 10 EGTA, 1 MgCl_2_, 1 CaCl_2_ (pH 7.2-7.3 with CsOH, 300-310 mOsm). Under such recording conditions, the calculated (pClamp v 8.2) and experimentally verified E_GABA_ and E_glutamate_ were -63 mV and 0mV respectively. Synaptic currents were filtered at 2 kHz using an 8 pole low-pass Bessel filter. The series resistance was between 8 and 20 MΩ with up to 80 % compensation. Only cells with a stable access resistance were used, and experiments were aborted if series resistance changes > 20% occurred.

#### Data analysis

All recordings were analyzed offline using the Strathclyde Electrophysiology Software (Electrophysiology Data Recorder [WinEDR] and Whole-Cell analysis Program [WinWCP]; courtesy of Dr J. Dempster, University of Strathclyde).

#### Analysis of sEPSCs & sIPSCs

Individual events were detected in WinEDR using an amplitude threshold detection algorithm (−4 pA threshold, 3 ms duration) and visually inspected for validity. Any noise that reached the threshold, or traces that contained multiple events were rejected from analysis. Accepted events (a minimum of 40 for each experimental condition) were digitally averaged. Synaptic currents (i.e. EPSCs and IPSCs) were analysed with regard to peak amplitude, rise time and decay kinetics. Events with rise times >1 ms (representing <1% of total) were discarded in order to eliminate from the analysis events subject to dendritic filtering. The decay phase of the digitally averaged event was then fitted with single exponential, [Y(t) = A*exp(-t/τ)] and bi-exponential functions, [Y(t) = A_1_*exp(-t/ τ_1_)+ A_2_*exp(-t/ τ_2_)]. An F test was then used to establish whether the decay was best described by a mono- or bi-exponential fit, as indicated by a decrease in the standard deviation of the residuals. The frequency of sEPSCs and sIPSCs was determined by counting the number of events in 20-second bins over a minimum of 2 separate 1-min. periods (first and last minute) of the recording. In WinEDR events were detected (offline) using a rate of rise time threshold that was specific for individual cells and that allows detection of the slowest events (∼40 pA/ms). Recordings were then visually inspected to ensure all events were included and any detected spurious noise removed. A mean frequency and inter-event interval were then calculated.

### Confocal Imaging of Microglial Density in the PVN

Perfused brains of CRH-Cre+/-::tdTomato+/-; CX3CR1-GFP+/+ mice (P10) were cryoprotected and frozen, then sectioned coronally into 25-µm-thick slices using a cryostat (1:4 series of the PVN). Unstained sections were mounted on gelatin-subbed slides and coverslipped with Vectashield containing DAPI. Confocal images of the PVN were collected with an LSM-510 confocal microscope (Zeiss) with an Apochromat ×20 oil objective. 11 z-stack images of 450 × 450 μm were taken at 1-μm intervals. Image frame was digitized at 12-bit using a 1024 × 1024 pixel frame size. An ROI was manually drawn around the perimeter of the CRH-tdTomato+ neurons of the PVN, then microglia number in this region was counted manually in FIJI, and microglial volume was automatically calculated using Imaris’ 3D reconstruction (Bitplane, Zurich, Switzerland).

### Two-photon Microscopy

#### Slice preparation

Acute PVN slices from P8 CRH-Cre+/-::tdTomato+/-; CX3CR1-GFP+/+ pups were prepared in the same manner as described above for Electrophysiology. Briefly, the brain was rapidly dissected and placed in ice cold (0-4°C), oxygenated (95% O_2_) aCSF, and 320-μm coronal hypothalamic slices were collected using a vibratome (Ted Pella, Redding, CA, USA). Slices were subsequently incubated for 30 min.-1 hr. at room temperature in a holding chamber containing oxygenated imaging media (50% MEM, 50% HBSS, 6 mg/mL D-glucose, and 25 mM HEPES; all from Invitrogen/ThermoFisher; adapted from 77 to be serum-free).

#### Imaging

Two-photon (2p) imaging of hypothalamic slices was performed using a multiphoton microscope built on an Olympus BX51 upright microscope frame, fitted with a motorized Z-Deck stage (Prior Scientific, Cambridge, UK) and Nikon 25x, water dipping objective (CFI75 Apo L, NA = 1.1, WD = 2.0 mm). 2P excitation was generated by a tunable femtosecond laser (Chameleon Ultra-II or Vision-II, Coherent, Santa Clara, CA, USA) set to 920 nm to excite EGFP and TdTomato as described previously^78^. Fluorescence emission was split by 484 nm and 538 nm dichroic mirrors into three non-descanned PMT detectors (R3896, Hamamatsu Photonics, Bridgewater, NJ, USA) to detect second-harmonic signal generated from collagen in blue; EGFP signal in green; and TdTomato signal in red. Supported brain slices were anchored in a RC-27L bath (Warner Instruments, Holliston, MA, USA) assembled into a custom-built heated imaging chamber maintained at 37°±0.5°C using continuous perfusion of oxygenated imaging media. 3D image stacks of x = 250 μm, y = 250 μm, and z = 56 μm (XYZ voxel size 0.244 µm x 0.244 µm x 4 µm) were acquired every 37 s interval using image acquisition software Slidebook (Intelligent Imaging Innovations, Denver, CO, USA), up to 50 min. to create a 4D data set. Imaris version 9.2.1 or later (Bitplane) was used for rendering and image processing for figures.

### Automated Python-based Algorithm for Analysis of Microglial Process Dynamics

#### Two-photon video pre-processing

All video files were converted from z-stacks to max projections using Slidebook software. The resulting videos had one 3-channel image per time point. All other processing code was written in Python. Videos were truncated to span 30 minutes of capture time for CTL vs. ELA experiments, or 50 min. for Gq-DREADD/CNO experiments, and only the green channel (containing microglia) was used for processing. To account for drift of tissue specimens during imaging, each frame in a given video was registered to the previous frame in the series using OpenCV’s^79^ *findTransformECC* and *warpAffine* functions.

Noise and point spread were reduced in each image using deconvolution (Fig. 2f). A measured point spread function for the microscope was unavailable, so one was generated with the FIJI (National Institutes of Health, Bethesda, MD, USA) plugin “PSF Generator” (http://bigwww.epfl.ch/algorithms/psfgenerator) using the following parameters:

|  |  |
| --- | --- |
| Model | Gibson & Lanni 3D Optical Model |
| Refractive index immersion | 1 |
| Refractive index sample | 1 |
| Working distance (ti) | 2,000 $\mu\text{m}$ |
| Particle position Z | 2,000 nm |
| Accuracy computation | Good |
| Wavelength | 920 nm |
| NA | 1.1 |
| Pixelsize XY | 244.140625 nm |
| Z-step | 4,000 nm |
| Size XYZ | 256 / 256 / 3 |
| Display | Linear / 32-bits / Grays |

The plugin generates a z-stack of 3 PSF images, and only the first slice of the z-stack was used for deconvolution of the 2D max projection image. The values in the PSF image were squared, as is necessary for two-photon images. Using the generated PSF image, each frame of the video was deconvolved using the Richardson-Lucy method (provided by the scikit-image80 function *skimage*.*restoration*.*richarson_lucy*).

After deconvolution, an optimal threshold for each video was determined manually in FIJI. Thresholds ranged from the top 10-15% of pixel values in the video and were chosen to remove background noise while maintaining microglia shape and branching. The thresholds were recorded but not applied to the videos at this point.

#### Soma finding

Microglial somas were located by thresholding the videos with a threshold that was 50% higher than the threshold determined by pre-processing. Erosion, dilation, and close operations were performed on the resulting binary image (using OpenCV’s^79^ *morphologyEx* function) to remove processes and fill in small gaps in the somas (Fig. 2f). Connected regions of non-zero pixels with an area greater than 281 pixels (approximately 17 microns^2^) were identified as somas.

Somas were tracked across frames in a video by centroid matching. Each soma in a frame is “matched” to the closest soma in the next frame if their centroids are less than 20 pixels apart. This process is repeated for each time point in a video to create a chain of somas across timepoints that belong to the same microglia. Occasionally, thresholding or processing criteria in previous steps that would result in a frame with a “missing” soma, where frames on either side have an identified soma in the same location but no soma was identified in the current frame. To account for this, somas in these “missing soma” frames were interpolated by overlapping (via binary “and” operation) the current thresholded image with the previous frame. Any connected region larger than 10 pixels that coincides with the location of a soma in the previous frame is labeled as a soma in the current frame.

#### Microglia tracing

Microglia structure was estimated by skeletonization (Fig. 2f). Deconvolved images were thresholded using the values determined during pre-processing, and connected regions of non-zero pixels with an area less than 20 pixels were removed from the thresholded image by setting their values to 0. The filtered binary image was skeletonized using the scikit-image80 function *skimage*.*morphology*.*skeletonize*.

To connect branches that became disconnected due to thresholding, each skeletonized image was turned into a graph using the minimum spanning tree (MST) algorithm (*scipy*.*sparse*.*csgraph*.*minimum_spanning_tree*^81^), which connects all points using the shortest path possible, with no cycles in the graph (Fig. 2f). Some modifications were made prior to running the algorithm. Briefly, a nearest-neighbor graph was generated for all non-zero points in the skeletonized image (using scikit-learn’s^82^ *sklearn*.*neighbors*.*kneighbors_graph*). The graph was modified to force all soma centroids to appear closer to the perimeter points of the soma than to any other point, which forced the MST algorithm to treat each soma as a tree center.

Additionally, the original pixel values of the deconvolved image were used as weights for the algorithm. The weight for each pair of points was calculated as the maximum (darkest) value between the points, which causes brighter pairs of pixels to be connected first. The end result was a graph describing how all skeletonized points connected to each other in the image. Prior to running MST, cycles in the original skeleton were identified. MST can break cycles in inconsistent places from image to image, so any “process tip” or branch that resulted from breaking a cycle was discarded from analysis. Tips that fell on the soma border were also discarded as they were artifacts of tracing outward from the soma center.

To identify individual microglia from the MST graph, all connections were broken between points that were greater than 50 pixels apart, as these were unlikely to belong to the same branch or microglia. If two or more somas were still connected to each other by a path, the path was broken at the longest gap between pixels in the skeletonized image. “Process tips” and branches resulting from these broken paths were discarded from analysis. This process resulted in individual graphs centered at each soma. Lastly, to reduce noise from skeletonization, individual branches that were less than 10 pixels long were pruned from the graph.

Similar to soma tracking, individual processes were tracked through frames in the video. Each process tip was “matched” to the closest process tip in the next frame, if it met three conditions. One, both tips must belong to the same microglia as identified by soma matching. Two, the tips must be less than 50 pixels away. Three, the tips must belong to the same branch. We determine this by tracing from each process tip to the nearest fork in its branch in each image. The branches must at some point in this path be less than 5 pixels away from each other to be considered the same. To reduce noise, only process tips that appear in 5 or more frames are considered for analysis.

#### Analysis

In addition to basic information about microglia location in each image in a video, information characterizing microglial process dynamics was also calculated. We define *tip velocity* between two frames as *d/t*, where *d* is equal to the Euclidean distance between matched process tips in two frames, and *t* is the time elapsed between frames.

#### Code availability

The Python tracking code is openly available on GitHub at https://github.com/jaclynrbeck/BaramLabMicrogliaTracking.

### Manual Kymograph Analysis of Microglial Process Dynamics

Pre-processing of 2-photon videos was similar to that described above for the automated method. The ProMoIJ plug-in for FIJI was used for drift correction/registration of the maximum projection, single-channel, time-lapse videos^83^, and the MultipleKymograph plug-in for FIJI (https://www.embl.de/eamnet/html/body_kymograph.html) was utilized to generate the kymographs themselves. In each video, 4 clearly delineated (located completely within the frame throughout the video), non-overlapping microglial cells were selected and cropped, and the brightest, most active process on each cell was selected for further analysis, resulting in 4 processes from 4 different microglia analyzed for each video. Next, an ROI was drawn as a line along the length of each microglial process, extending from the base where the process connects to the cell body, to slightly past the process tip. The kymograph analysis measured the total distance travelled by the tip of the microglial process (Fig. 2d), which was averaged across the 4 analyzed processes to attain one average value for each animal.

### *Ex Vivo* Chemogenetic Activation 2p Imaging Experiment

The preparation of acute hypothalamic slices from P8 CX3CR1-Cre+::Gq-DREADD+ mice was conducted as described above. Rather than EGFP, mCitrine-labeled Gq-DREADDs labeled microglia in this strain and were used to track microglial process dynamics. A baseline period of 10 min. was imaged using the same parameters as before. At 10 min., the perfusion tubing was switched to imaging media containing 10 μM CNO (CAS# 34233-69-7, MH# C-929, National Institutes of Mental Health, Bethesda, MD, USA) in 0.05% dimethyl sulfoxide (DMSO; cat# D8418-100, Millipore Sigma) or 0.05% DMSO alone (Vehicle) for 40 more min. of imaging to determine the effect of CNO vs. Vehicle. 2p videos were analyzed using the automated Python-based algorithm and manual kymograph method as described above.

### Microglial Synapse Engulfment Analysis

Coronal sections of the PVN from P8 CRH-Cre+/-::tdTomato+/-; CX3CR1-GFP+/+ mice were subjected to vGlut2 IHC using a similar procedure as described above for the synapse IHC. Briefly, after several washes in PBS containing 0.3% Triton X-100 (PBS-T, pH 7.4), sections were blocked with 5% normal donkey serum (Jackson ImmunoResearch) for 1 hr. to prevent non-specific binding. Sections were then incubated overnight at 4°C with guinea pig anti-vGlut2 antiserum (1:12,000, cat# AB2251-1, Millipore Sigma) in PBS-T containing 2% normal donkey serum. The next morning, sections were rinsed in PBS-T (3 × 5 min.), and then incubated with donkey-anti-guinea pig IgG-647 (1:1,000; cat# 706-605-148, Jackson Immunoresearch) for 3 hr. at room temperature. Sections were washed several times in PBS-T, mounted on gelatin-coated slides, and coverslipped with Vectashield mounting medium with DAPI (Vector Laboratories, H-1200). Confocal images of the mpd PVN were collected with an LSM-510 confocal microscope (Zeiss) with an Apochromat ×63 oil objective. 11 z-stack images of 142.86 × 142.86 μm were taken at 1-μm intervals. Image frame was digitized at 12-bit using a 1024 ×1024 pixel frame size. An ROI was manually drawn around the perimeter of the CRH-tdTomato+ neurons of the mpd PVN, and microglial volume within the ROI was automatically calculated using Imaris’ 3D reconstruction. Excitatory vGlut2+ synaptic puncta located inside of the 3D microglial volume were automatically identified and counted using Imaris’ spot detection function.

### Microglial Ultrastructure Analysis

Two PFA/acrolein-perfused brain sections containing the PVN were selected from a P8 CRH-Cre+/-::tdTomato+/- male mouse (PVN sections were selected based on the presence of densely packed CRH-tdTomato+ neurons around the 3rd ventricle). Sections were washed in PBS, then quenched 10 min. in 0.3% H_2_O_2_ in PBS and permeabilized 30 min. in 0.1% NaBH_4_ in PBS. Sections were first incubated 1 hr. at room temperature in blocking solution (10% fetal bovine serum, 3% bovine serum albumin, 0.01% Triton X-100 in [50 mM] tris-buffered saline [TBS]). Afterwards, sections were incubated overnight at 4°C with rabbit anti-IBA1 polyclonal primary antibody (1:1,000; cat# 019-19741, FUJIFILM Wako Chemical, Osaka, Japan) in blocking solution. The following day, antibody was washed off, and the sections were incubated with biotinylated goat anti-rabbit polyclonal secondary antibody (cat#111-066-046, Jackson ImmunoResearch,) in TBS for 1.5 hr., followed by avidin-biotin complex solution (1:1:100 in TBS; cat# PK-6100, Vector Laboratories) for 1 hr. at room temperature. The staining was revealed in 0.05% diaminobenzidine (DAB; cat# D5905-50TAB, Millipore Sigma) with 0.015% H_2_O_2_ in TBS for 4.5 min. at room temperature.

The immunostained sections were next post-fixed flat in osmium-thiocarbohydrazide-osmium for scanning electron microscopy (SEM). In particular, sections were incubated in 3% ferrocyanide (cat# PFC232.250, BioShop, Burlington, ON, Canada) diluted in water combined (1:1) with 4% aqueous osmium tetroxide (cat#19170, Electron Microscopy Sciences, Hatfield, PA, USA) for 1 hr. in 1% thiocarbohydrazide diluted in water (cat# 2231-57-4, Electron Microscopy Sciences) for 20 min., in 2% osmium tetroxide diluted in water, then dehydrated in ascending concentration of ethanol (2 × 35%, 50%, 70%, 80%, 90%, 3 × 100%) followed by propylene oxide (3×) for 5 min. each. After post-fixation, tissues were embedded in Durcupan ACM resin (cat# 44611-44614, Millipore Sigma) for 24 hr. and carefully placed between two ACLAR® embedding films (cat# 50425-25, Electron Microscopy Sciences), and the resin was let to polymerize at 55°C for 72 hr. Regions of selection—PVN—were excised from the embedded sections on ACLAR® sheets and re-embedded on top of a resin block for ultrathin sectioning (Ultracut UC7 ultramicrotome, Leica Microsystems). Ultrathin sections (∼75 nm thickness) were collected and placed on a silicon nitride chip and glued on specimen mounts for SEM. Representative microglial cell bodies and processes in the PVN were imaged at 5 nm of resolution using a Crossbeam 540 field emission SEM with a Gemini column (Zeiss).

### Mer IHC

Coronal sections of the PVN from P8 CRH-Cre+/-::tdTomato+/- male mice were subjected to Mer IHC using a similar procedure as described above for the synapse IHC. Briefly, after several washes in PBS containing 0.3% Triton X-100 (PBS-T, pH 7.4), free-floating sections were incubated in 0.5% Triton X-100 in PBS for 10 minutes, followed by 2 additional washes in PBS-T. Sections were blocked in 5% normal donkey serum (NDS) and 5% normal goat serum (NGS) in PBS-T, followed by incubation overnight at 4°C with rat anti-Mer (1:1000, eBioscience, cat# 14-5751-82) and rabbit anti-P2RY12 (1:2000, AnaSpec, cat# AS-55043A) in 2% NGS and 2% NDS in PBS-T. Following several washes in PBS-T, sections were incubated with goat anti-rat IgG conjugated to Alexa-Fluor 488 (1:500, Invitrogen Life Technologies, cat# A11055) and donkey anti-rabbit IgG conjugated to Alexa-Fluor 647 (1:500, Jackson ImmunoResearch Inc., cat# 711-605-152) in 2% NDS and 2% NGS in PBS-T for 3 hr. at room temperature. Sections were washed several times in PBS-T, mounted on gelatin-coated slides, and coverslipped with Vectashield mounting medium with DAPI (Vector Laboratories, H-1200). Confocal images of the PVN were collected with an LSM-510 confocal microscope (Zeiss) with an Apochromat ×20 oil objective. 11 z-stack images of 450 × 450 μm were taken at 1-μm intervals. Image frame was digitized at 12-bit using a 1024 × 1024 pixel frame size. An ROI was manually drawn around the perimeter of the CRH-tdTomato+ neurons of the PVN, then Mer volume and microglial volume (based on P2RY12+ immunoreactivity) were automatically calculated using Imaris’ 3D reconstruction.

### Mer Inhibitor Treatment of Organotypic PVN Slice Cultures

P6-7 male CRH-Cre+/-::tdTomato+/- mouse pups were rapidly decapitated, brains were removed from the skull, and hypothalamic blocks were dissected and cut into 350-μm coronal sections on a McIlwain tissue chopper. To create organotypic PVN cultures, up to 6 slices were collected posterior to the anterior commissure. These slices were explanted onto Millicell cell culture inserts (pore size 0.4 µm, diameter 30 mm, Merck Millipore Inc., cat# PICMORG50). Membrane inserts were placed into a six-well plate with 1 mL of culture medium. Culture medium consisted of 52% modified Eagle’s medium [MEM; cat# 11700, Invitrogen], 25% Hanks Balanced Salt Solution [HBSS; cat# 24020, Invitrogen], 20% Heat Inactivated Horse Serum (added post-filtration) supplemented with 3mM L-Glutamine (Gibco, cat# 25030081), 25mM D-Glucose (Sigma, cat# G7528), 1.9mM NaHCO3 (ThermoFisher, cat# 25080094), 12.5mM HEPES (Gibco, cat# 15630080), 0.6mM L-Ascorbic acid (Sigma-Aldrich, cat# A4403), 1µg/mL Insulin (Sigma-Aldrich, cat# I0516) and 25µg/mL containing penicillin-streptomycin (Gibco, cat# 15140122). Slices were cultured at 37°C in 5% CO_2_ enriched air for 48 hr., at which point the medium was refreshed. After an additional 48 hours, the medium was refreshed with medium without antibiotics. 6 days after initial culture, each membrane was washed twice with 2 mL of antibiotic-free, serum-free medium (97% MEM supplemented with 3mM L-Glutamine, 10mM D-Glucose,1.9mM NaHCO3, 12.5mM HEPES, 0.6mM L-Ascorbic acid, 1µg/mL Insulin) and treatment with 20nM of a small-molecule Mer inhibitor (UNC2025, Selleck Chemicals, cat# S7576) or sterile culture-grade water as vehicle was commenced. Media containing either vehicle or Mer inhibitor was refreshed 12 hr. later. After an additional 4 hr., cultures were fixed in 4% paraformaldehyde in 0.1 M phosphate-buffer (PB) on ice for 30 min. rinsed in PB, and cryoprotected in a 25% sucrose solution for 4-6 hr.

#### Organotypic PVN slice cultures IHC and imaging

Following cryoprotection, each slice confirmed to contain PVN was individually frozen on dry ice while still attached to the membrane. Slices were sectioned to 14-µm thickness using a Leica CM1900 cryostat and were immediately mounted onto gelatin-coated slides. Slides were stored at -20°C until further processing. Upon thawing, slides were rinsed in PBS-T, followed by treatment in 0.3% H2O2 in 0.01 M PBS for 20 min. Sections were blocked in PBS-T containing 5% NDS and then incubated overnight at 4°C with rabbit anti-PSD95 (1:1000, Invitrogen/ThermoFisher) and guinea pig anti-vGlut2 (1:10,000, Millipore Sigma) in PBS-T containing 2% NDS. Sections were washed in PBS-T and then incubated in donkey anti-guinea pig IgG conjugated to Alexa-Fluor 647 (1:500, Jackson ImmunoResearch) and donkey anti-rabbit pig IgG conjugated to Alexa-Fluor 488 (1:500, ThermoFisher) in PBS-T containing 2% NDS for 3 hr. at room temperature. Sections were washed with PBS-T and coverslipped with Vectashield mounting medium with DAPI (Vector Laboratories). Confocal images of the mpd PVN were collected with an LSM-510 confocal microscope (Zeiss) with an Apochromat ×63 oil objective. 11 z-stack images of 142.86 × 142.86 μm were taken at 1-μm intervals. Image frame was digitized at 12-bit using a 1024 ×1024 pixel frame size. CRH neuronal volume was automatically calculated using Imaris’ 3D reconstruction function. Excitatory synapses onto CRH+ neurons were identified as colocalized puncta of vGlut2+PSD95 within the CRH-tdTomato+ volume using Imaris’ colocalization function (threshold=1.0).

### *In Vivo* Chemogenetic Activation Experiments

CX3CR1-Cre+::Gq-DREADD+ mice were crossed with Gq-DREADD+ mice to generate ∼50% of pups expressing excitatory Gq-DREADDs specifically in their microglia (Fig. 4a; Fig. S5), and the rest served as littermate controls. On P3, small, sustained-release CNO- or placebo-containing pellets (CNO: cat# X-999, 0.025 mg/pellet, 8-day release; Placebo: cat# C-111; Innovative Research of America, Sarasota, FL, USA) were inserted under the skin (s.c.) of male pups to obviate the stress of daily injections. Litters were culled to 6 pups maximum, with at least 1 female. Also on P3, the litters were randomized to CTL or ELA rearing for a week (Fig. 4b; Fig. S1a). One cohort of mice were transcardially perfused on P10 (as described above) to measure the number of excitatory synapses onto PVN mpd neurons after CTL vs. ELA rearing. A separate cohort was transferred to standard cages at P10 and grown up until adulthood, then transcardially perfused and adrenal glands collected and weighed as a measure of lifetime chronic stress. In another cohort, adult mice were subjected to multiple acute stresses (MAS)^59,60^ in order to assess their response to stress in adulthood.

### IHC for excitatory synaptic markers and confocal imaging in P10 Gq-DREADD brains

Perfused brains of P10 CX3CR1-Cre+::Gq-DREADD+ mice were post-fixed, cryoprotected, and cryosectioned as described above. PVN-containing sections were subjected to IHC protocols similar to those described above. Briefly, after several washes with PBS containing 0.3% Triton X-100 (PBS-T, pH 7.4), sections were blocked with 5% normal donkey serum (Jackson ImmunoResearch) for 1 hr. to prevent non-specific binding. Sections were then incubated overnight at 4°C with rabbit anti-PSD95 antiserum (1:1,000, Invitrogen/ThermoFisher) and guinea pig anti-vGlut2 antiserum (1:12,000, Millipore) in PBS-T containing 2% normal donkey serum. The next morning, sections were rinsed in PBS-T (3 × 5 min.), and then incubated with donkey-anti-rabbit IgG-568 (1:1,000; cat# A10042, ThermoFisher) and donkey-anti-guinea pig IgG-647 (1:1,000; Jackson ImmunoResearch) for 3 hr. at room temperature. After washing (3 × 5 min), sections were mounted onto gelatin-coated slides and coverslipped with Vectashield containing DAPI (Vector Laboratories). Confocal images of the mpd PVN were collected using an LSM-780 confocal microscope (Zeiss) with an Apochromat ×63 oil objective. 11 z-stack images of 142.86 × 142.86 μm were taken at 1-μm intervals. Image frame was digitized at 12-bit using a 1024 × 1024 pixel frame size. An ROI was manually drawn around the perimeter of the densely packed DAPI+ nuclei of the mpd PVN. Excitatory synapses in this region were identified as colocalized puncta of vGlut2+PSD95 using Imaris’ colocalization function.

### Multiple acute stresses (MAS) paradigm in adulthood

The MAS paradigm consists of restraint, bright light, unpredictable loud noise, physical jostling and awareness of peer discomfort (as described in ^60^), and has been previously shown to elicit a robust stress response greater than a single stressor alone^84^. Briefly, adult male mice were put in a restrainer made from a 50-ml plastic tube (Corning, Corning, NY, USA), placed next to unfamiliar conspecifics (social stress) on a laboratory shaker, and jostled in a brightly lit room with loud rap music playing (dB level= 95) for 1 hr. Small blood samples (∼100 μL) were collected from the facial vein of each mouse at baseline, 30 min. after MAS initiation, and 60 min. after MAS initiation, and serum was collected after clotting for ∼30 min. at room temperature and centrifugation at 1,200x*g* for 15 min. Serum was stored at -20°C until later assayed for ACTH and CORT using commercially available enzyme-linked immunosorbent assay kits according to the manufacturers’ instructions (ACTH kit: cat# EK-001-21, Phoenix Pharmaceuticals, Burlingame, CA; CORT kit: cat# 501320, Cayman Chemical, Ann Arbor, MI).

### Statistical Analysis

Differences between CTL and ELA groups were assessed using unpaired t-tests, with Welch’s correction for unequal variance as necessary. One-sample t-tests were used for comparing CNO to Vehicle (i.e., 100%) in the ex vivo chemogenetics experiment. 2-way ANOVA was used to analyze the effects of ELA and Mer inhibition separately and in interaction (ELA X Drug), followed by Tukey’s post hoc tests. Differences after ELA and the in vivo chemogenetics intervention were analyzed by one-way ANOVA or Welch’s ANOVA (corrected for unequal variance) as necessary, followed by the appropriate post hoc tests (specified for each test in Results section). Significance levels were set at 0.05, and data are presented as mean ± SEM. Grubbs’ test was used to remove statistical outliers from the data. Statistical analyses were performed using GraphPad Prism 9.0 software (GraphPad, San Diego, CA, USA). All experiments were assessed blindly without prior knowledge of the experimental group.

