## Supplemental Material for "Impaired developmental microglial pruning of excitatory synapses on CRH-expressing hypothalamic neurons exacerbates stress responses throughout life"

**Supplementary Files: Bolton et al.**

| | sEPSCs CTL (n=9) | ELA (n=13) | sIPSCs CTL (n=9) | ELA (n=15) |
|---|---|---|---|---|
| **Peak ampl. (pA)** | -31 ± 3 | -30 ± 2 | 43 ± 7 | 44 ± 3 |
| **Rise time (ms)** | 0.4 ± 0.1 | 0.4 ± 0.1 | 0.5 ± 0.1 | 0.4 ± 0.1 |
| **Decay τ (ms)** | 1.5 ± 0.1 | 1.5 ± 0.1 | 7.7 ± 0.4 | 6.8 ± 0.5 |
| **Freq (Hz)** | 1.6 ± 0.3 | 3.0 ± 0.5* | 1.5 ± 0.4 | 1.6 ± 0.3 |

**Supplementary Table S1.** Summary of the properties of spontaneous excitatory and inhibitory postsynaptic currents recorded from mediodorsal parvocellular (mpd) paraventricular hypothalamic nucleus (PVN) neurons derived from CTL and ELA mice. Data are mean ± SEM; *$p$<0.05.

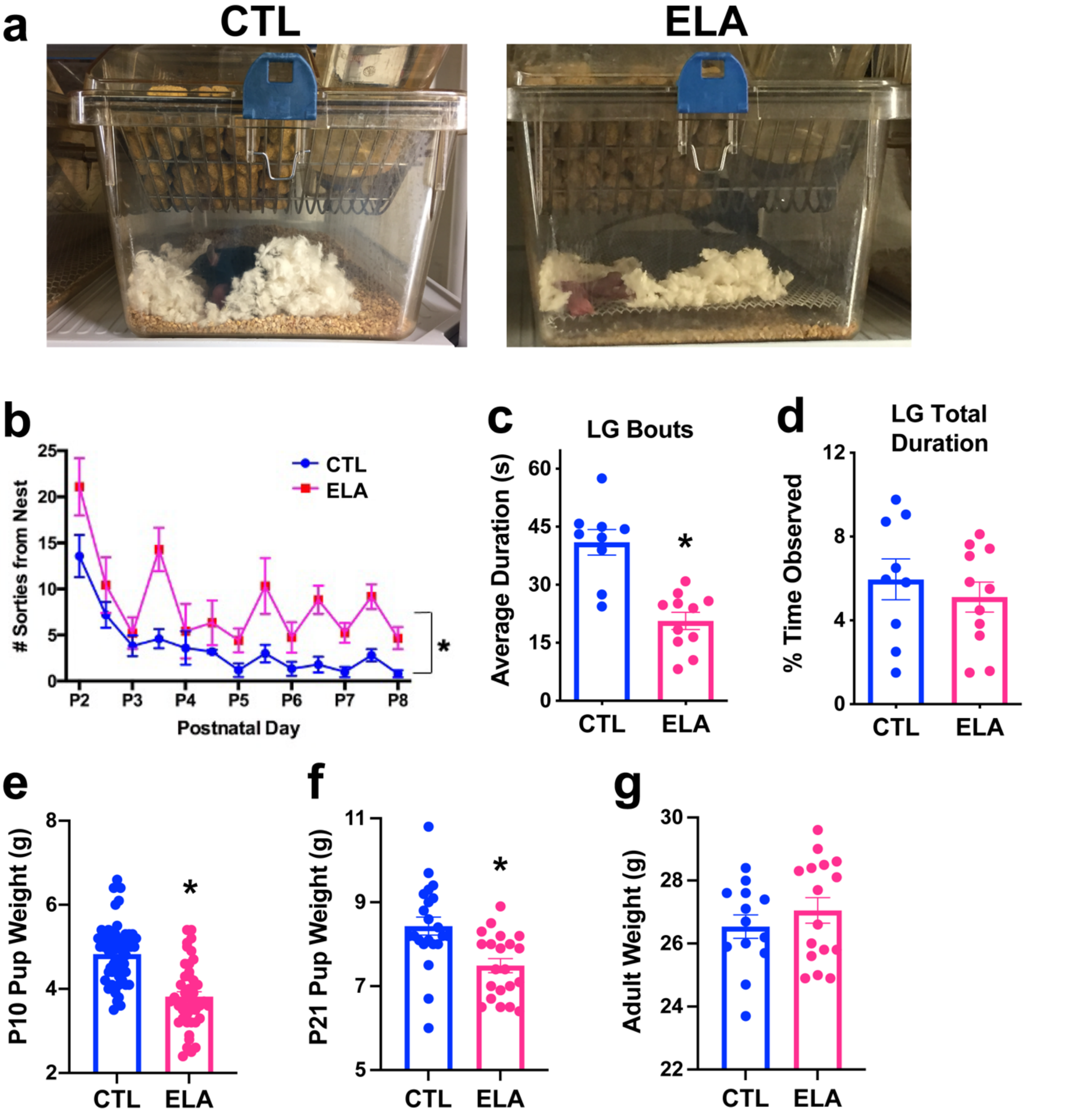


**Supplementary Figure S1: Limited bedding and nesting model of early-life adversity (ELA).** a. Representative photographs of the standard cage environment (CTL; left) and LBN cage environment (right) employed to generate ELA. b. Dams rearing their litters in LBN cages made significantly more sorties (i.e., exits) from the nest during the ELA period of postnatal days (P)2-8 than CTL dams ($F_{1,19}$=25.15, p<0.0001; 2-way repeated measures ANOVA). c. ELA dams exhibited fragmentation of their maternal care behaviors, particularly licking and grooming (LG) of their pups, as measured by the average duration of a LG bout during P2-8. d. ELA does not alter the total duration of LG, as measured by the percent of time observed during P2-8, indicating that ELA alters the pattern, rather than the overall quantity, of maternal care. e. ELA provokes diminished pup body weights by the end of the ELA period on the morning of P10 ($t_{101}$=7.05, p<0.0001; unpaired t-test). f. The ELA-induced decrease in body weight is maintained until weaning at P21. g. However, body weights of ELA offspring normalize by adulthood (P60). Data are mean ± SEM; **p*<0.05.

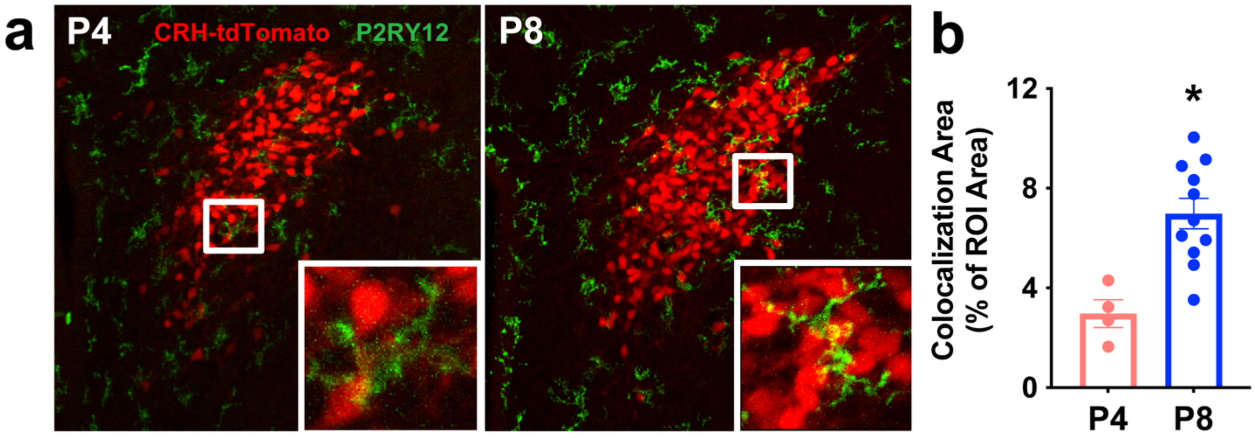


**Supplementary Figure S2: Microglial colocalization with CRH+ neurons in the paraventricular hypothalamic nucleus (PVN) across the first postnatal week.**

a. Representative confocal images of P4 (left) and P8 (right) PVN CRH-expressing neurons (tdTomato-labeled, red) and microglia (labeled with P2RY12-488, green). b. The amount of colocalization or overlap between microglia and CRH+ neurons (as a % of ROI area, defined by the clustering of CRH+ neurons) increases from P4 to P8 ($t_{13}=3.71$, $p=0.003$; unpaired t-test), suggesting that microglia interact with CRH+ neurons more closely at the end of the first postnatal week than at the beginning. Data are mean ± SEM; $*p<0.05$.

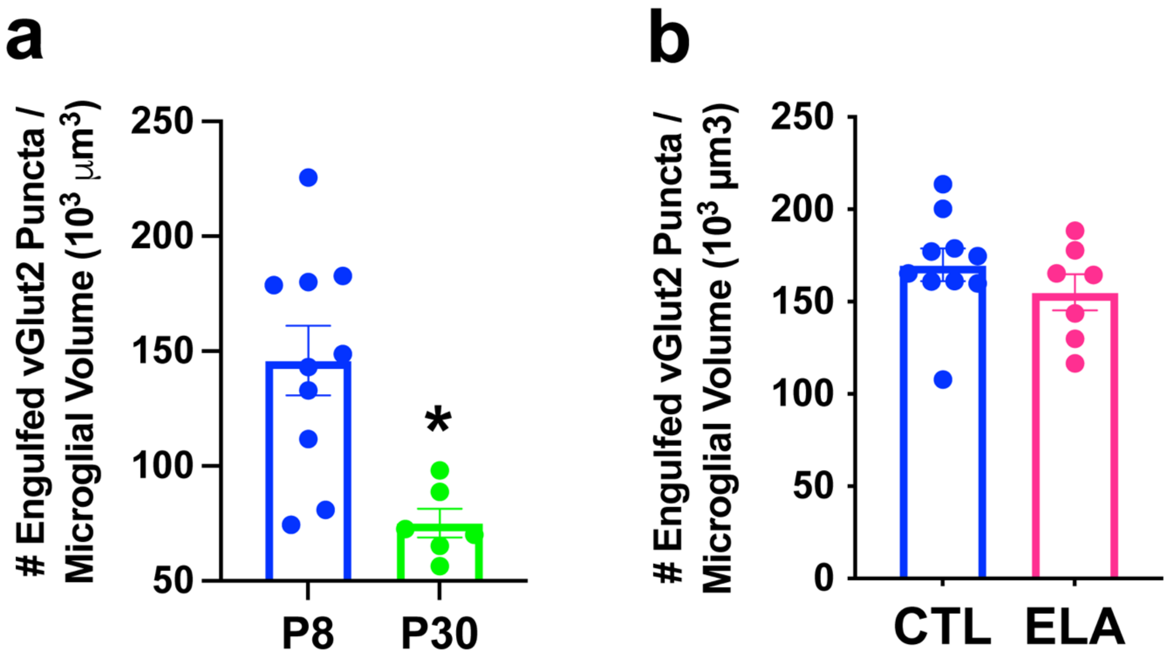


**Supplementary Figure S3: Properties of microglial synapse engulfment in the immature mediodorsal parvocellular (mpd) paraventricular hypothalamic nucleus (PVN).**
a. Postnatal day (P)8 is a period of active synapse engulfment, with levels of engulfed vGlut2+ synaptic puncta twice as high as at P30 in CTL mice ($t_{11.75}$=4.31, p=0.001; Welch's t-test). b. Microglia that were on the periphery of the PVN and not directly contacting CRH+ neurons did not exhibit the ELA-induced decrease in synapse engulfment (p>0.2) observed in microglia that were abutting CRH+ neurons in P8 ELA mice. Data are mean ± SEM; *$p$<0.05.

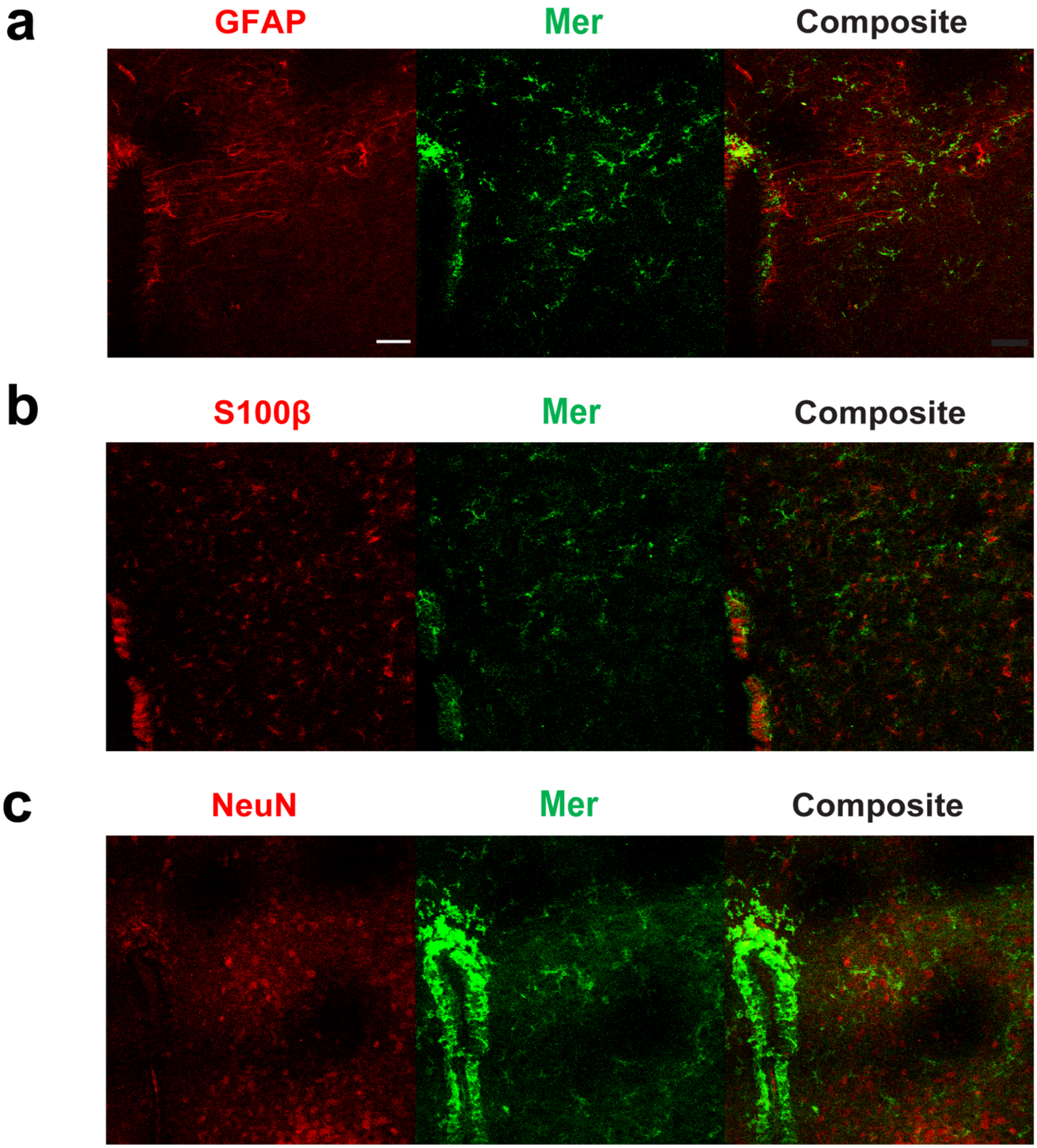


**Supplementary Figure S4: Mer cell type-specific expression in the immature paraventricular hypothalamic nucleus (PVN).**
a. Representative confocal image of Mer immunoreactivity (green) not colocalizing with GFAP immunoreactivity (red; labels astrocytes) in the P8 PVN. Scale bar=50 μm. b. Representative confocal image of Mer immunoreactivity (green) not overlapping with S100β immunoreactivity (red; labels astrocytes) in the P8 hypothalamus. c. Representative confocal image of Mer immunoreactivity (green) distinct from NeuN immunoreactivity (red; labels neurons) in the P8 hypothalamus.

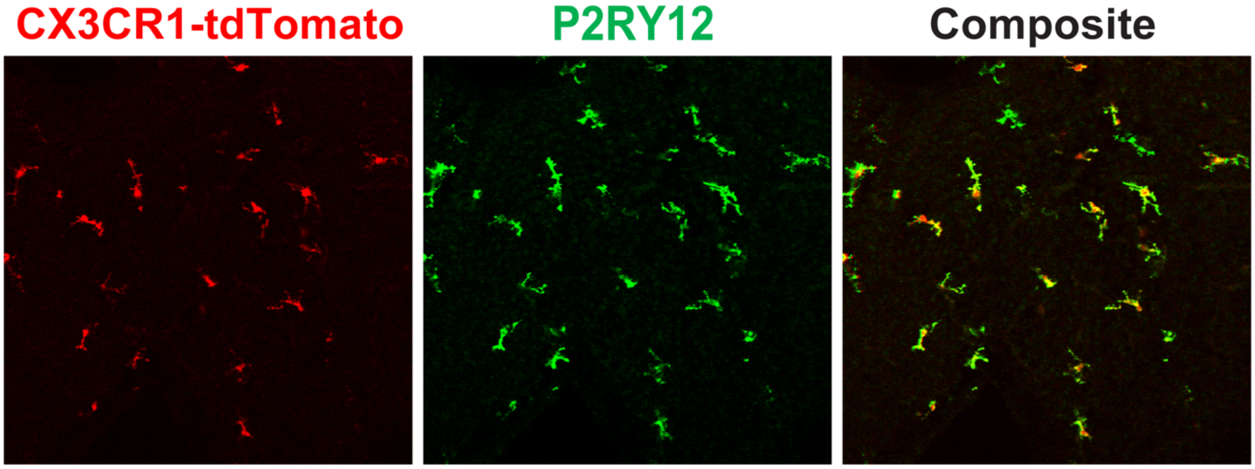


**Supplementary Figure S5: Specificity of Cre expression to microglia in CX3CR1-Cre mice.** Representative confocal image of hypothalamic microglia (labeled with P2RY12 immunostaining; green) expressing Cre (tdTomato; red; overlap shown in yellow in composite image) in a CX3CR1-Cre+::tdTomato+ mouse soon after birth, on postnatal day (P)0.

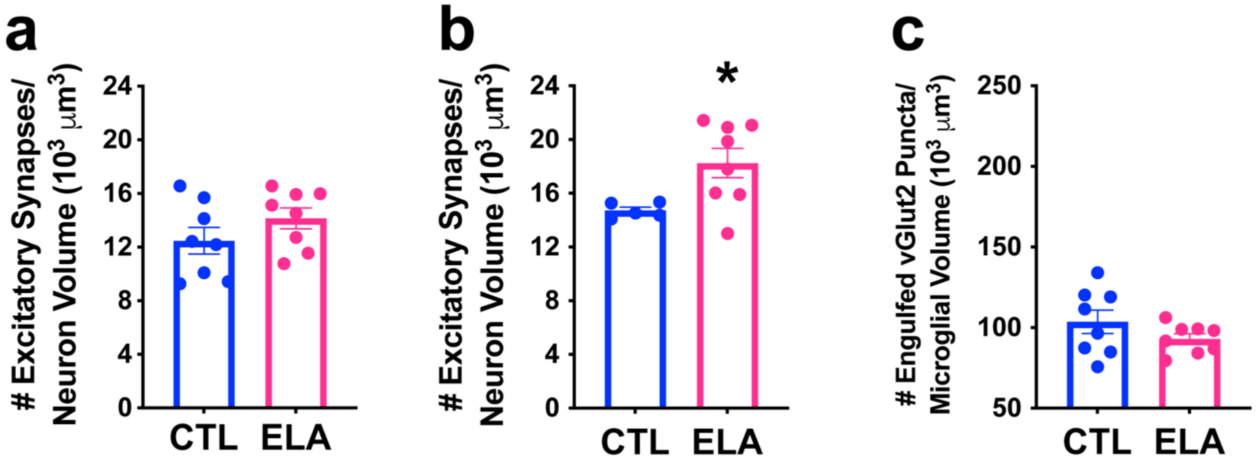


**Supplementary Figure S6: Neonatal female mice do not exhibit the same microglial mechanism of ELA-induced synapse excess as their male littermates.**

a. Female ELA pups do not exhibit a significant augmentation of excitatory synapses on CRH+ neurons in the mediodorsal parvocellular (mpd) paraventricular hypothalamic nucleus (PVN) at the end of the ELA period on postnatal day (P)10 ($t_{14}$=1.33, p=0.2; unpaired t-test). b. ELA does increase excitatory synapse number on CRH+ neurons in female offspring by P24-5 ($t_{7.73}$=3.17, p=0.01; Welch's t-test). c. ELA does not provoke a significant decrease in microglial synapse engulfment in P8 female mpd PVN, ($t_{9.2}$=1.68, p=0.1; Welch's t-test), unlike in ELA males. Data are mean ± SEM; *$p$<0.05.

**Movie S1. Visualization of microglial process dynamics.** 2-photon time-lapse movie from postnatal day (P)8 PVN with fluorescent protein-labeled microglia (green) and CRH+ neurons (red). Maximum intensity projections showing microglia engaging multiple CRH cells. 3D image frames were acquired at 45-sec intervals; scale bar=10 μm; time is shown as hr:min:sec; playback speed is15 fps.
